# Allele-specific rescue of neurexin behavioral phenotypes by monoamine-targeting compounds

**DOI:** 10.64898/2026.02.06.703315

**Authors:** William R. Smisko, William R. Haury, Michele Perni, Rebecca Kalik, Jhelaine Palo, Brandon L. Bastien, Jadwiga N. Bilchak, Raheem Khadour, Thomas A. Jongens, Matthew S. Kayser, Michael P. Hart

## Abstract

Neurexins are synaptic adhesion molecules associated with neurologic changes in humans, including neurodevelopmental delay, autism, schizophrenia, Tourette syndrome, and seizures. The *NRXN1* gene produces >100 protein isoforms through alternative promoters and extensive splicing, which are differentially impacted by *NRXN1* variants found in patients. Yet pharmacologic targeting of *NRXN1* isoforms or deletions has not been comprehensively studied. Here, we developed a behavioral screening approach in *C. elegans* to identify small molecule compounds that modify the decreased activity levels caused by isoform-specific deletions of neurexin(*nrx-1*). Screening 190 compounds, we discovered that monoamine-targeting compounds differentially improve behavioral phenotypes depending on which *nrx-1* isoforms are disrupted. Broad modulation of monoamine signaling, or antagonism of specific serotonin receptors, are required to increase the activity of both alleles tested. The FDA-approved atypical antipsychotic olanzapine was the sole validated compound achieving Z-scores >2 in both screens, which notably also rescued behavioral phenotypes of *C. elegans* harboring a conserved autism-associated *NRXN1* missense variant (L18Q/L16Q) identified in human patients. In *Drosophila Nrx-1* mutants, olanzapine uniquely and significantly improved activity deficits and extended survival, demonstrating evolutionary conservation of our findings. Multi-behavior testing revealed pharmacological specificity: olanzapine improved both activity and social feeding phenotypes of *nrx-1* alleles, while asenapine maleate improved activity, but worsened social feeding, indicating distinct impacts across behavioral domains. Our findings establish monoamine modulation as a conserved compensatory mechanism for neurexin loss, identify olanzapine as a lead compound for targeting neurexin loss, and demonstrate that allele stratification and pharmacogenomic approaches are needed for precision intervention in behavioral conditions.

Neurexins are synaptic adhesion molecules implicated in autism, schizophrenia, and neurodevelopmental disorders. *NRXN1* produces over 100 isoforms differentially affected by patient variants, yet pharmacologic targeting has not been systematically studied. We developed a *C. elegans* behavioral screen to identify compounds rescuing activity deficits caused by isoform-specific deletions. Screening 190 compounds, we discovered that monoamine-targeting drugs differentially improved behavioral phenotypes depending on which isoforms were disrupted. The FDA-approved antipsychotic olanzapine uniquely achieved robust rescue (Z-scores >2) across all genetic backgrounds tested, including a conserved autism-associated *NRXN1* missense variant (L18Q/L16Q) identified in patients. Olanzapine’s efficacy was conserved in *Drosophila Nrx-1* mutants, improving activity and extending survival. Multi-behavior testing revealed pharmacological specificity: olanzapine rescued both activity and social feeding phenotypes, while asenapine differentially affected behavioral domains. Our findings establish monoamine modulation as a compensatory mechanism for neurexin loss and identify olanzapine as a therapeutic lead for precision intervention in neurexin-associated disorders.

## INTRODUCTION

Animal behavior is regulated by circuits that are collections of neurons connected by synapses. Synaptic adhesion molecules organize the development, function, and plasticity of these circuits, and disruption of these genes can profoundly alter behaviors. Neurexins (*NRXN1-3*) are well-characterized synaptic adhesion molecules associated with neurodevelopmental delay, autism, schizophrenia, Tourette syndrome, seizures, and other behaviors^1–6^. In animal models, neurexin disruption affects sleep, social behaviors, learning, and motor function^7–13^. The three mammalian neurexin genes are exceptionally complex, producing 2-3 major isoforms from alternative promoters (α, β, and γ (*NRXN1*-specific)) with extensive splicing, potentially generating hundreds to thousands of unique neurexin proteins^14–16^. These isoforms exhibit neuron and synapse-specific localization and function^14,17–19^, where individual neurons express transcripts encoding dozens of NRXN1 isoforms^19^. Deletions and variants in human *NRXN1* have isoform-specific impacts^20–22,2,19,4^ and can result in treatment-resistant anxiety, depression, and seizures, highlighting a need for neurexin genotype-focused interventions.

Despite the importance of neurexins to neurodevelopment and their association with multiple neurodevelopmental and neuropsychiatric conditions, we lack a comprehensive understanding of how they alter behaviors, which has limited effective interventions. Identifying pharmacological modifiers of neurexin-related behavioral phenotypes could reveal mechanisms underlying behavioral changes, compensatory pathways, and potential therapeutic targets. However, fundamental questions remain: Can synaptic adhesion molecules, typically considered structural organizers, or defects arising from their loss, be targeted pharmacologically? Should interventions target the proteins directly or focus on compensatory circuit and signaling pathways? Can compounds broadly or specifically target the wide array of behavioral changes associated with *NRXN1* disruption? One report screened for small molecule inhibitors of a NRXN1-LRRTM physical interaction, but follow-up and functional characterization of identified inhibitors was limited^23^. An alternative approach using behavioral phenotypes to identify functional modifiers of neurexin isoforms and circuit function could reveal direct and indirect modifiers. Despite the clinical importance of neurexins and synaptic adhesion molecules, few or no compounds targeting these proteins or impacted pathways exist, leaving open the opportunity to target them as dynamic regulators of behavior for intervention^24^.

We recently reported multiple foraging behavioral phenotypes in *C. elegans* with loss of function alleles in synaptic adhesion molecules, including neurexins (*nrx-1*), neuroligins (*nlg-1*), and CNTNAPs (*nlr-1*)^25–27^. *C. elegans* respond to acute food loss with sustained increases in activity lasting over 8 hours, a compensatory foraging response regulated by sensation and integration of food cues by sensory neurons and monoamine interneurons^28–32^. We found that loss of *nrx-1* reduces the behavioral response to food deprivation, with isoform-specific alleles showing distinct temporal effects^25^. The *nrx-1* γ isoform regulates behavioral initiation through octopamine signaling from RIC interneurons, while the *nrx-1* α isoform regulates behavioral maintenance through presynaptic structural and other changes^25^. A related gene, *nlr-1,* functions in dopamine neurons to regulate this behavior, and a small pharmacological screen identified dopamine receptor-specific modifiers^27^, demonstrating the feasibility to uncover genotype specific circuit mechanisms and targets. *C. elegans* behavioral phenotypes are ideal for modifier screening due to their small and well-defined nervous system, conserved neuronal signaling pathways, rapid life cycle, and tractable genetics and behaviors. However, to establish translational relevance, findings need to be validated through additional mechanisms, which could include testing in models of patient-identified variants or alternative model organisms that recapitulate neurexin dysfunction.

Here, we screened 190 FDA-approved small-molecule compounds for their ability to improve activity levels in food deprived *C. elegans* harboring isoform-specific *nrx-1* alleles. We identified compounds that selectively increase activity in *nrx-1* null mutants and, in a second screen, discovered shared and allele-specific compounds in mutants lacking only the α *nrx-1* isoform. The top modifier compounds predominantly target monoamine signaling, showing distinct temporal dynamics and magnitude of effects across control and *nrx-1* genotypes. Behavioral phenotypes resulting from disruption of all (α and γ) or only α *nrx-1* isoforms were rescued by modifiers targeting broad monoamine receptors and/or antagonizing specific serotonin receptors. However, loss of only *nrx-1* α-isoform was uniquely rescued by modulation of adrenergic/octopaminergic signaling, while loss of γ isoform required additional antagonism of dopamine signaling. We validated compounds in a second behavioral phenotype, social feeding, revealing behavior-specific effects: asenapine maleate rescued food deprivation responses, but worsened social feeding phenotypes, while olanzapine (an FDA-approved atypical antipsychotic) rescued both behaviors. We also find that multiple monoamine compounds improve comparable behavioral phenotypes resulting from insertion of a conserved patient reported *NRXN1* variant into *nrx-1* (L18Q/L16Q). Finally, we tested the top candidate compounds in *Drosophila Nrx-1* mutants, which have reduced activity levels and reduced survival, and found that olanzapine significantly increased their activity and probability of survival. Our results identify the first pharmacological modifiers of neurexin behavioral phenotypes, reveal allele- and behavior-specific modifiers and disrupted pathways, and demonstrate cross-species potential for compounds targeting monoamine pathways.

## METHODS

### C. elegans husbandry

*C. elegans* were grown on plates with Nematode Growth Media (NGM) agar seeded with *Escherichia coli* OP50 bacteria as a food source and raised at 20°C. Unless otherwise noted, the N2 Bristol strain was used as the control strain, and assays were performed on day 1 adult animals except when indicated otherwise. Strains are listed in **Supplemental Table 5**.

### Pharmacologic compound behavioral analysis in WormCamp setups

An optical 96-well plate (Optically Clear Tissue Culture Plate, 96 Well, 190 uM clear base, black frame, Azenta Life Sciences) was used as a **Worm C**ollective **A**ctivity **M**onitoring **P**latform (WormCamp)^33^. The WormCamp measures the collective activity of a population of animals in each well^27^. We screened using the Tocriscreen FDA-approved drugs library (Tocris brand at Bio-Techne), purchased at 10 mM in DMSO (**Supplemental Table 1**). These compounds were added to wells at concentrations indicated and analyzed for impact on behavior in the WormCamp setup. Strains were synchronized using standard alkaline hypochlorite solution methods (bleaching) approximately forty-eight hours before setting up the assay allowing for the assay to be run with animals as day one adults. The optical 96-well plate was prepared one hour before strain populations reached adulthood. The inner sixty wells of the 96-well plate were filled with 74 μL of M9 buffer using a multichannel pipette. The surrounding edge wells were not used to avoid edge effects from the WormWatcher camera system due to light refraction. 1 μL of compound from the library or DMSO vehicle was pipetted into each well. For compound screening, each plate included two replicate wells per compound, and six replicates for vehicle alone (0.1 mM final concentration in well). Two or three independent replicate plates were run for each compound and each strain. The plate was left covered with aluminum foil on an orbital shaker at low speed for one hour to equilibrate.

Fifteen minutes prior to the end of the mixing period, the synchronized animals were washed off agar plates with food into a 15 mL conical tube. Once all animals were combined into the conical tube, they were centrifuged at low speed for two minutes. The supernatant with food was aspirated off to ensure food-deprived conditions, and the animals were resuspended in 3 mL of M9 buffer. Centrifugation and aspiration were repeated as needed if the supernatant contained visible food. To achieve ∼30 animals per well, animals were pipetted up and down to distribute evenly, and then 25 μL was pipetted onto a microscope slide. Animals were counted under a dissecting scope, and M9 buffer was adjusted to dilute the concentration to 30 ± 5 animals / 25 μL. The concentration was verified at least three additional times to ensure uniformity. Animals were transferred into a 25 mL Aspir-8™ Reservoir and put on an orbital shaker at high speed to ensure even suspension and distribution. To load the 96-well plate, a multichannel pipette was used to pipette 25 μL of the diluted animal suspension with the orbital shaker adjusted between pipetting to ensure suspension in solution. After all wells were filled with 25 μL (final volume of 100 μL per well), they were visually analyzed to note wells with very low numbers of animals (<10 or fewer animals per well), and any such wells were noted and removed from analysis.

For single worm WormCamp assays the above protocol was modified. Fifty of N2 Bristol controls and the *nrx-1* strains were picked onto new seeded plates as late L4’s. Animals were transferred to the 96 well plate at day 1 adulthood. The optical 96 well plate was prepared the same as mentioned before, except 98 μL of M9 buffer was added to each well initially. The top and bottom rows were avoided, and an alternating quadrant layout was used to further mitigate positional effects. Quadrants were broken up into 6×3 well sections (12 replicates per strain per plate) with 1% DMSO vehicle, and two drug concentrations (0.125 mM and .0625 mM), each row containing one condition. N2 Bristol controls and the *nrx-1* allele each had two alternating quadrants. To transfer animals, day 1 adults were picked into unseeded NGM plates (one for each strain) with 3 mL of M9 buffer. Plates were mixed to wash animals of food. Each animal was pipetted individually in 1 μL of M9 into their respective well. Each compound had 2-3 replicate plates for *nrx-1(wy778)*, *nrx-1(nu485),* and *nrx-1(sy869)* mutants, each being directly compared to N2 Bristol controls.

The loaded 96-well WormCamp plate was then placed in the WormWatcher imaging platform (Tau Scientific Instruments, West Berlin, NJ, USA). WormWatchers captured images of the WormCamp for 9 hours. 12 images were captured per session (session is equal to 1 minute), with sessions sequentially imaged every 10 minutes, and 6 sessions per hour. Images were processed using a previously reported frame subtraction method^34^ and analysis software created by Tau Scientific Instruments, that segments individual wells using pixel subtraction on consecutive images to calculate the activity by change in pixels. To normalize activity to zero, 5 hours of activity for each well was divided by the average activity of all DMSO vehicle alone wells on the same replicate plate for those 5 hours.

### Aggregate feeding behavior assay

Standard 6-well plates with 6 mL of NGM (nematode growth medium) in each well were seeded with 75 μL of OP50 bacterial culture that contained either 0.4% vehicle (DMSO) or 0.1 mM compound in DMSO (or concentration indicated in DMSO). Plates were seeded with vehicle or small molecule compound solution, left covered and shielded from light to dry, approximately 20 hours before running the assay. The following day, L4 mutant and control *npr-1(ad609)* animals were collected. Each replicate had *npr-1(ad609)* aggregating control animals with vehicle control and with two small molecules or two concentrations of the same small molecule. This was repeated for *npr-1(ad609);nrx-1(wy778)*, *npr-1(ad609);nrx-1(nu485)*, and *npr-1(ad609);nrx-1(sy869)* mutant animals. 2-3 independent replicates were performed per genotype and compound concentration. Fifty animals of the indicated genotype were placed onto the center of each well in the center of the bacterial lawn, to ensure minimize disturbance of the lawn to avoid creation of denser areas of food. After the animals were loaded into wells, 10x Tween was thinly coated onto the lid to avoid condensation during imaging. Plates were placed into a WormWatcher imaging platform, and 12 images were taken over 5-second intervals in the first minute of each hour for 20 hours. The number of aggregating *C. elegans* per well was manually counted from blinded images, where any animal in contact with two or more other animals was considered aggregating, as previously described^26^. In cases where the number of aggregating animals could not be clearly counted, single animals were counted and subtracted from 50 total to obtain a count of aggregating *C. elegans*. Data shown is from hour 15 after experimental set-up, therefore representing day 1 adult animals.

### *Drosophila* genetics and husbandry

Drosophila strains containing the Nrx-1^273^ and Nrx-1^241^ alleles are previously described^8,35^. Flies were outcrossed in the w1118(iso31Bw-) background for six generations. Flies were maintained on a standard cornmeal molasses medium obtained from Lab Express (Fly Food R, recipe available at http://lab-express.com/DIS58.pdf) on a 12-hour:12-hour light:dark (LD) cycle at 25°C. All fly experiments utilized trans-heterozygous mutants, carrying one *Nrx-1^273^* allele and one *Nrx-1^241^* allele.

### *Drosophila* drug preparation

Glass tubes with drugs were prepared as previously described^36^. Drugs or DMSO vehicle stocks were diluted to the final concentration (0.025-0.25mM in 1% DMSO or 1% DMSO only respectively) in a 5% sucrose/2% agar solution.

### *Drosophila* locomotor activity analysis and survival

Trans-heterozygous *Nrx-1* mutant male flies and their respective controls (iso31Bw-) were collected 1 to 2 days after eclosion and aged for 3 days in group housing on standard food at 25°C on a 12-hour:12-hour LD cycle. Flies were then anesthetized on CO_2_ pads (Genesee Scientific, catalog no. 59-114) and loaded into glass tubes containing 5% sucrose and 2% agar and Drugs or DMSO as described above. All experiments were loaded between ZT5-ZT7. All data collection began at ZT12 at least 5 hours following CO2 anesthesia. Locomotor activity was monitored using the DAM5H multibeam system (Trikinetics, Waltham, MA). Activity was measured in 12h bins^37^. Data was processed with a custom R script using Rethomics package^38^. Activity index was calculated as the average number of beam breaks per minute of wake time. Activity and survival data was processed with a custom R script using Rethomics package^38^

### Statistics and reproducibility

Statistical analyses were performed and data were plotted in GraphPad Prism 10 or RStudio. Unbiased computer vision analysis determined the activity values for WormCamp. For library screen data, mean activity was calculated for each well and compound wells were normalized to the DMSO alone wells from the same plate (and of the same genotype). For single animal WormCamps run with controls, mean activity was calculated for each well and normalized to the control N2 DMSO wells from the same plate. For WormCamp behavioral experiments, data were plotted with each dot representing activity of an individual well across independent replicates. To identify statistically significant screening hits, we calculated the Z-scores for each compound in both screens according to the formula Z = (X - μ) / σ, where X is the compound’s activity, μ is the mean activity across all compounds, and σ is the standard deviation. Z-scores indicate how many standard deviations a compound’s effect lies from the screen mean, with |Z| > 2 corresponding to p < 0.023 (approximately 2.3% false positive rate). This threshold was used to identify high-confidence hits for validation and further characterization. For validation and characterization experiments, compound wells were compared to DMSO alone control wells using a one-way ANOVA with Tukey’s Post-Hoc test applied. For social feeding behavioral experiments, the hour 15 counts of aggregating animals were plotted for each genotype. Each data point represents an individual well of a 6-well plate. Plots include the standard error of mean (SEM). To compare aggregation behavior levels across genotypes, a one-way ANOVA was performed with a Tukey’s Post-Hoc test applied. For *Drosophila* activity experiments means for each tube were calculated for the timepoints indicated; each dot represents an individual animal. A one-way ANOVA was performed with Dunnet’s test to compare activity across all genotypes and treatments, p-values are indicated on each graph. For *Drosophila* survival experiments, statistical analysis, a Logrank (Mantel-Cox) test was used (Control vs *Nrx-1*, **** p<0001; Control vs Control + 0.025mM, NS p=0.2756; Control vs Control + 0.25mM, NS p=0.3118; *Nrx-1* vs *Nrx-1* 0.025mM, NS p=0.4117; *Nrx-1* vs *Nrx-1* 0.25mM *** p=0.0002).

## RESULTS

### Screening for small molecule modifiers of a nrx-1 behavioral phenotype

We previously determined that both the long α and short γ isoforms of *nrx-1* are required for adult *C. elegans* to respond to food deprivation with increased activity levels, which is likely a compensatory foraging behavior and mild stress response. We found that exposure to exogenous octopamine in the presence of food increased activity of control animals, and animals with loss of the α isoform, but loss of both *nrx-1* isoforms (α and γ) reduced this effect^25^. Modification of this behavior with octopamine led us to develop an assay to screen for other small molecules that increase the activity of *nrx-1* mutant animals in the absence of food. In part with the goal to identify signaling and neuronal pathways impacted by, or that can compensate for, loss of *nrx-1* isoforms. To easily deliver small molecules to *C. elegans* we replicated the previous behavioral assay but transitioned from animals crawling on solid agar (individual animals per well on 48-well WorMotel) to populations of animals swimming in liquid (∼30 animals per well in a 96-well plate (WormCamp)^27^. Using this setup, we found that populations of day 1 adult animals lacking *nrx-*1 α and γ isoforms (*wy778*) or lacking just the *nrx-1* α isoform (*nu485*) had reduced swimming activity compared to control animals in the absence of food (**Figure 1A-B**).

**Figure 1.**
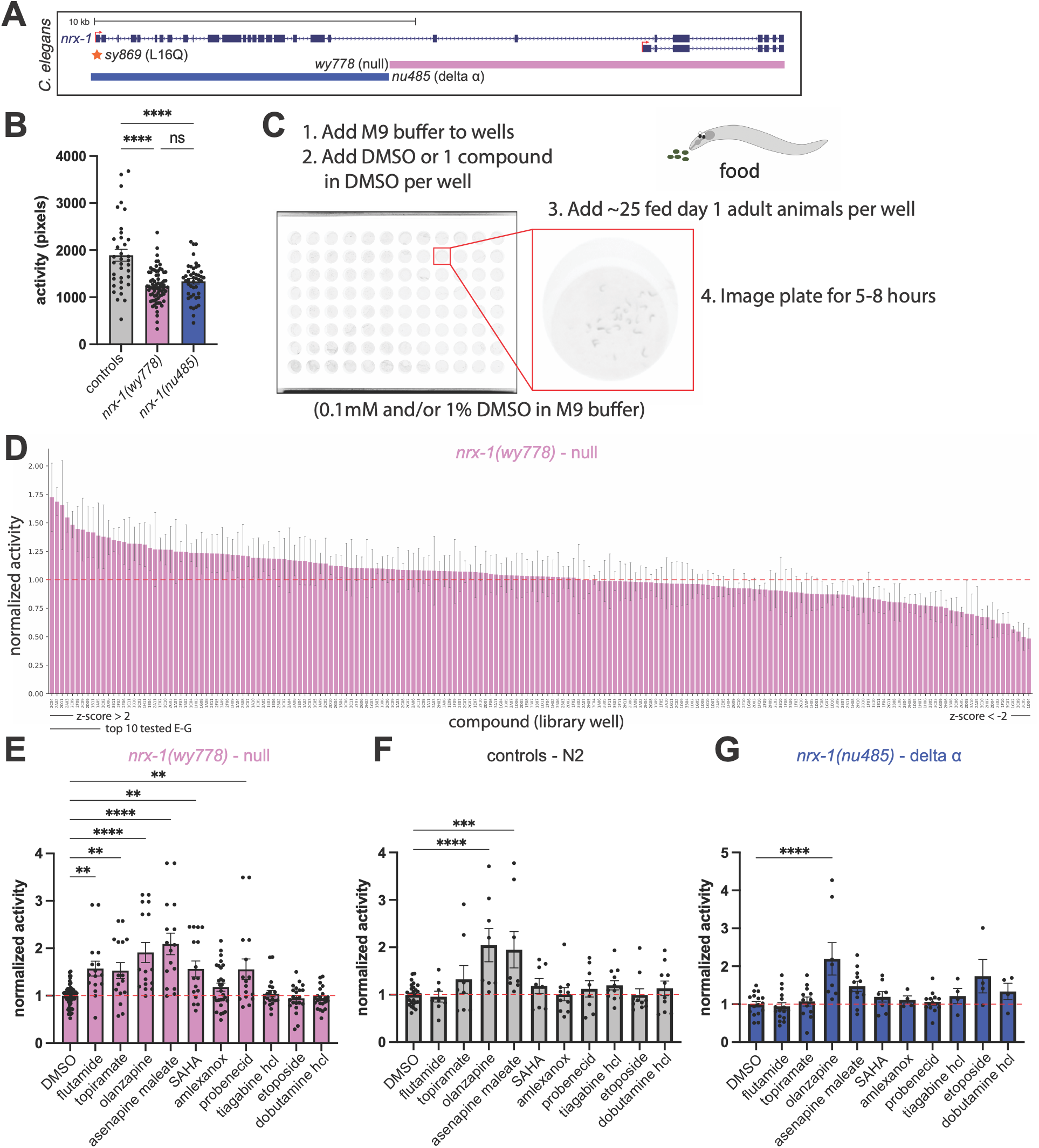
Screen and identification of compounds that increase activity of *nrx-1* null animals without food. **A)** Diagram showing *nrx-1* genomic locus, with examples of two major isoforms (α and γ) and alleles used in this study (*wy778* deletion in pink, *nu485* deletion in blue, and *sy869* missense variant in orange). **B)** Quantification of activity levels of day 1 adult control, *nrx-1(wy778)*, or *nrx-1(nu485)* without food. **C)** Cartoon schematic of compound screening and behavioral assay. **D)** Activity levels of ∼25 animals per well swimming in liquid with no food and 0.1 mM of each compound (190 from TocriScreen library) in 1% DMSO normalized to animals (4 well for each compound across 2 independent plates) in DMSO vehicle alone (1%)(compounds ordered from highest to lowest average normalized activity, black lines under x-axis indicate the compounds with z-scores >2 or <-2 and top 10 compounds further validated in panels below). Secondary test of top 10 compounds from same library with activity levels in for each compound normalized to DMSO vehicle controls to validate in (**E**) *nrx-1(wy778)* animals, and test for first time in (**F**) N2 control, and (**G**) *nrx-1(nu485)* animals.

We screened 190 small molecule compounds in the WormCamp for impact on activity levels of day 1 adult animals lacking both isoforms of *nrx-1(wy778)*. Each compound (in DMSO) was tested at 0.1 mM across multiple wells in 2 independent plates tested on separate days (1% total DMSO)(**Figure 1C**). The impact of compounds on activity levels of *nrx-1(wy778)* animals was calculated by normalizing 5 hours of activity of each well to multiple wells of *nrx-1(wy778)* animals in only 1% DMSO as a vehicle alone control. Of the 190 compounds in the library, we found that a subset increased activity levels of *nrx-1(wy778)* animals compared to vehicle alone (**Figure 1D**). The top 10 compounds that increased mean activity levels were **flutamide, topiramate, olanzapine, asenapine maleate, SAHA**, amlexanox, probenecid, tiagabine hydrochloride, etoposide, and dobutamine hydrochloride (bold had Z scores above 2, p < 0.023)(**Figure 1D** and **Supplemental Table 1**), which include compounds targeting diverse signaling pathways described in more detail below (monoamine receptors, androgen receptor, GluK1 and carbonic anhydrase, histone deacetylases, multidrug transporters, and others). We also identified compounds that lowered activity levels compared to vehicle alone, including vilazodone hydrochloride, dofetilide, sorafenib, FTY 720, delavirdine mesylate, axitinib, metformin hydrochloride, roflumilast, **felodipine**, **lapatinib**, **clotrimazole**, and **isradipine** (bold had Z scores below -2, p < 0.023)(**Figure 1D** and **Supplemental Table 1**). Of these, two are calcium channel blockers, which could inhibit movement and activity of animals (isradipine and felodipine), and three are inhibitors of growth factor receptors (lapatinib, axitinib, and sorafenib). There is also a single monoamine targeting compound, the potent 5-HT1A partial agonist vilazodone hydrochloride.

To validate and more carefully quantify the screening results focusing on compounds that increased activity, we tested the top 10 compounds using the same behavioral setup and concentration (0.1mM with total of 1% DMSO) in 8 or more wells across 2 or more replicate plates (**Figure 1E**). We found that 6 of the top 10 compounds significantly increased activity levels of *nrx-1(wy778)* animals over 5 hours compared to DMSO vehicle alone, including the top 5 compounds and the 7^th^ compound (**Figure 1E**). Interestingly, we found that the impact of the 3^rd^ and 4^th^ compounds (olanzapine and asenapine maleate) resulted in the largest increases in activity (**Figure 1E**), and both compounds broadly target monoamine signaling. In control animals, we found that only 2 of the top 10 compounds significantly increased activity levels (olanzapine and asenapine maleate)(**Figure 1F**). We also tested the impact of the top 10 compounds on activity of animals lacking only the α isoform (*nu485*)(comparing to the respective genotypes in vehicle alone, which does not allow comparison of magnitude of impact across genotypes). Only 1 of the top 10 significantly increased activity levels in *nrx-1(nu845)*(olanzapine)(**Figure 1G**). These results validate and confirm our behavioral screening results in *nrx-1(wy778)*, and suggest we can identify shared, allele-, and genotype-specific behavioral modifier compounds.

### Validation of compounds identified in screening of null nrx-1 animals

To further characterize modifier compounds with the largest impact across diverse targets, we tested multiple concentrations of asenapine maleate, topiramate, and flutamide acquired from different sources (**Supplementary Table 2**). For each of these three modifiers, we also tested two compounds with similar predicted targets (**Supplementary Table 2**). For asenapine maleate, a serotonin antagonist that also targets dopamine, histamine, and adrenergic receptors, we tested the impact of pimethixene maleate and perospirone, both serotonin and dopamine antagonists. All three of these compounds increased activity of *nrx-1(wy778)* animals without food, with the highest concentrations having largest impact, although perospirone also increased activity of *nrx-1(wy778)* animals at even lower concentrations (**Figure 2A-C**). These 3 compounds had less impact on activity levels of control animals, where asenapine lowered activity at 0.1 mM (**Figure 2D**). Interestingly, the two asenapine similar compounds significantly increased activity in control N2 animals at 0.03 mM (**Figure 2E-F**). We again found that asenapine did not increase activity of *nrx-1(nu485)* animals without food, and interestingly also significantly lowered activity at 0.1 mM (**Figure 2G**). The asenapine similar compounds also had no consistent impact on activity levels of *nrx-1(nu485)* animals, although pimethixene maleate significantly lowered activity at 0.01 mM (**Figure 2H-I**). Together, these results confirm the impact of top monoamine compounds at the concentration screened, and the finding that targeting monoamine modulation has genotype-specific impacts^25^.

**Figure 2.**
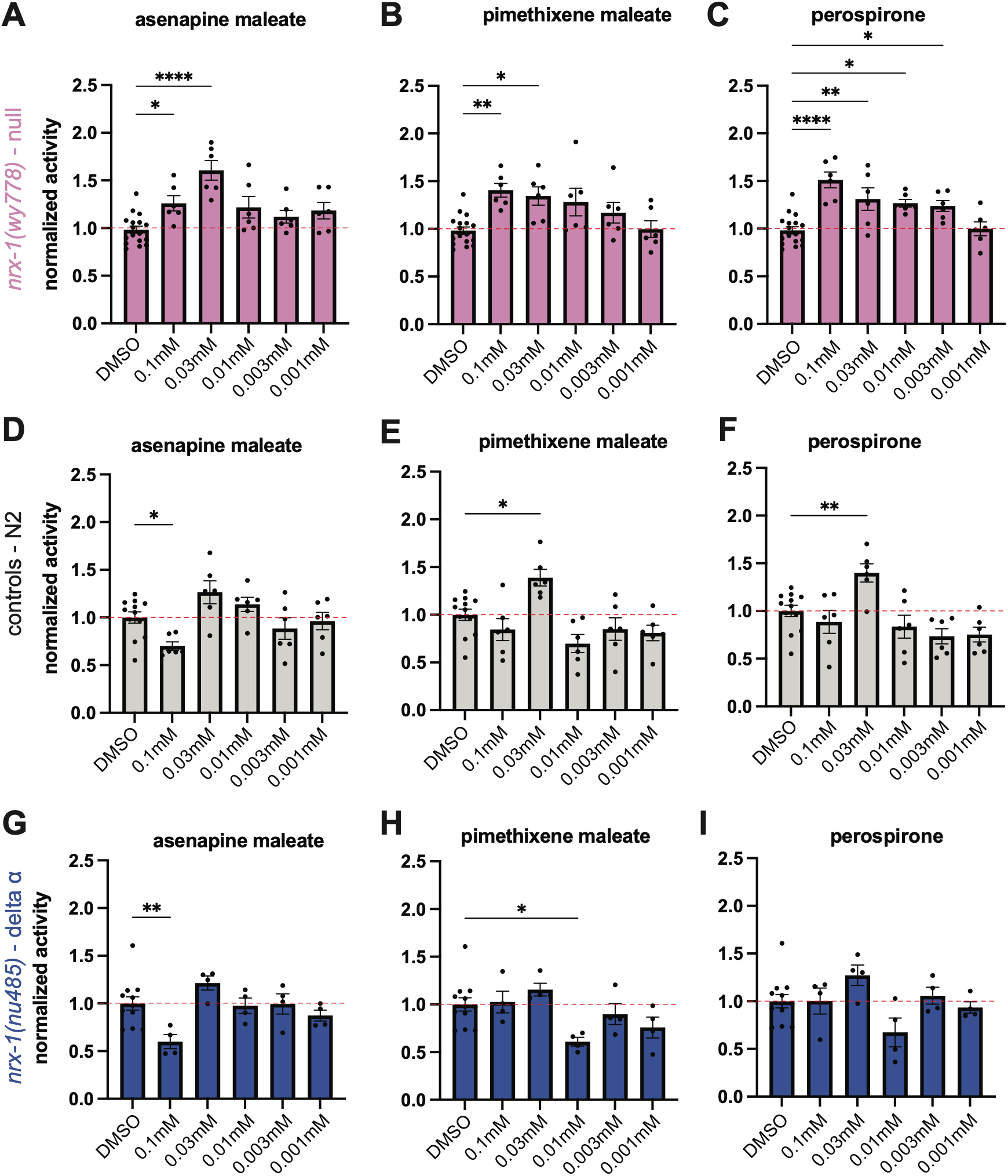
Validating compounds identified in *nrx-1(wy778)* null screen. Activity levels of ∼30 animals per well swimming in liquid with no food and a range of concentrations of each compound (asenapine maleate, pimethixene maleate, or perospirone) and 1% DMSO normalized to wells of animals in DMSO vehicle alone (1%) for (**A-C**) *nrx-1(wy778)*(pink), (**D-F**) controls (grey), and (**G-I**) *nrx-1(nu485)*(blue).

For topiramate, which inhibits voltage-gated sodium channel and calcium channels, AMPA/kainate receptors, and carbonic anhydrase, we selected acetazolamide (carbonic anhydrase inhibitor) and CNQX (AMPA and kainate receptor antagonist) for our similar predicted target compounds (**Supplemental Table 2**). In *nrx-1(wy778)* and *nrx-1(nu485)* animals, 0.1 mM of topiramate slightly increased activity but was not significant, while lower concentrations had no impact or subtle decreases on activity (**Supplemental Figure 1A-B**). We observed no impact of topiramate on control animals (**Supplemental Figure 1C**). The similar predicted target compounds did not have any significant impact on the activity of *nrx-1(wy778)* animals and only subtle impacts on control and *nrx-1(nu485)* activity levels (**Supplemental Figure 1A-C**). Exposure to 0.3 mM flutamide slightly increased activity of *nrx-1(wy778)*, without reaching significance, and had no impact on activity levels of controls and *nrx-1(nu485)* animals (**Supplemental Figure 1A-C**). The two similar predicted target compounds for flutamide had either no impact on activity levels of all genotypes, or a significant decrease in activity, with ostarine decreasing activity levels of all genotypes at high concentration(s), and spironolactone slightly decreasing *nrx-1(wy778)* activity at a medium/low concentration (**Supplemental Figure 1A-C**). Together, this confirms genotype-specific modifier compounds, with differences in the magnitude of impact arising when testing compounds from different sources. With no consistent impact of the top hit similar compounds of topiramate and flutamide, we did not identify specific signaling impacted, which may reflect compound or source specific differences.

### Screening for modifiers of behavioral phenotype of animals lacking only α isoform

The validation and characterization of the top compounds from the *nrx-1(wy778)* modifier screen, identified differential impacts on controls and animals lacking the long α isoform of *nrx-1* (*nu485*). Therefore, we screened the same 190 compound library to identify modifiers of the similar behavioral phenotype of *nrx-1(nu485)* animals. Based on the results from the *nrx-1(wy778)* screen we tested each compound at 0.1mM in 1% DMSO for 5 hours without food across multiple wells in at least 2 independent plates tested on separate days (1% total DMSO) (**Figure 1C**). We normalized activity of each well to average activity of the wells of *nrx-1(nu485*) animals with DMSO vehicle alone. We identified several compounds that significantly modified activity levels of *nrx-1(nu485)* animals (**Figure 3A**). The top 10 compounds that increased activity of *nrx-1(nu485)* animals were **dihydroergotamine mesylate, olanzapine, probenecid, dexmedetomidine hydrochloride**, fenofibrate, valsartan, mirtazapine, abacavir hemisulfate, bosutinib, and zolmitriptan (bold have Z scores above 2, p < 0.023)(**Supplemental Table 3**). When comparing the results from both *nrx-1* screens using an xy plot from both screens, there are modifiers with shared and *nrx-1* allele-specific impacts (**Figure 3A&B, Supplemental Table 4**). A Pearson correlation between screens indicated a weak but significant correlation (*p* = 2e-04 and *R* = 0.27)(**Figure 3B**), suggesting some similarity, but also differences, in modifier profiles between alleles, despite being deletion alleles of a single gene. The *nrx-1(nu485)* screen validated the previous observation of the shared impact of olanzapine on the two *nrx-1* alleles, with a Z score greater than 2 in both screens making it a statistical outlier (**Figure 1 G&E, Supplemental Tables 3&4**). The compounds that lowered activity in the *nrx-1(nu485)* screen included some shared and distinct compounds with the *nrx-1(wy778)* screen. For example, lapatinib, axitinib, and vilazodone hydrochloride lowered activity in both *nrx-1* alleles. Uniquely lowering activity of *nrx-1(nu485)* were additional monoamine targeting compounds, pimozide and rizatriptan benzoate.

**Figure 3.**
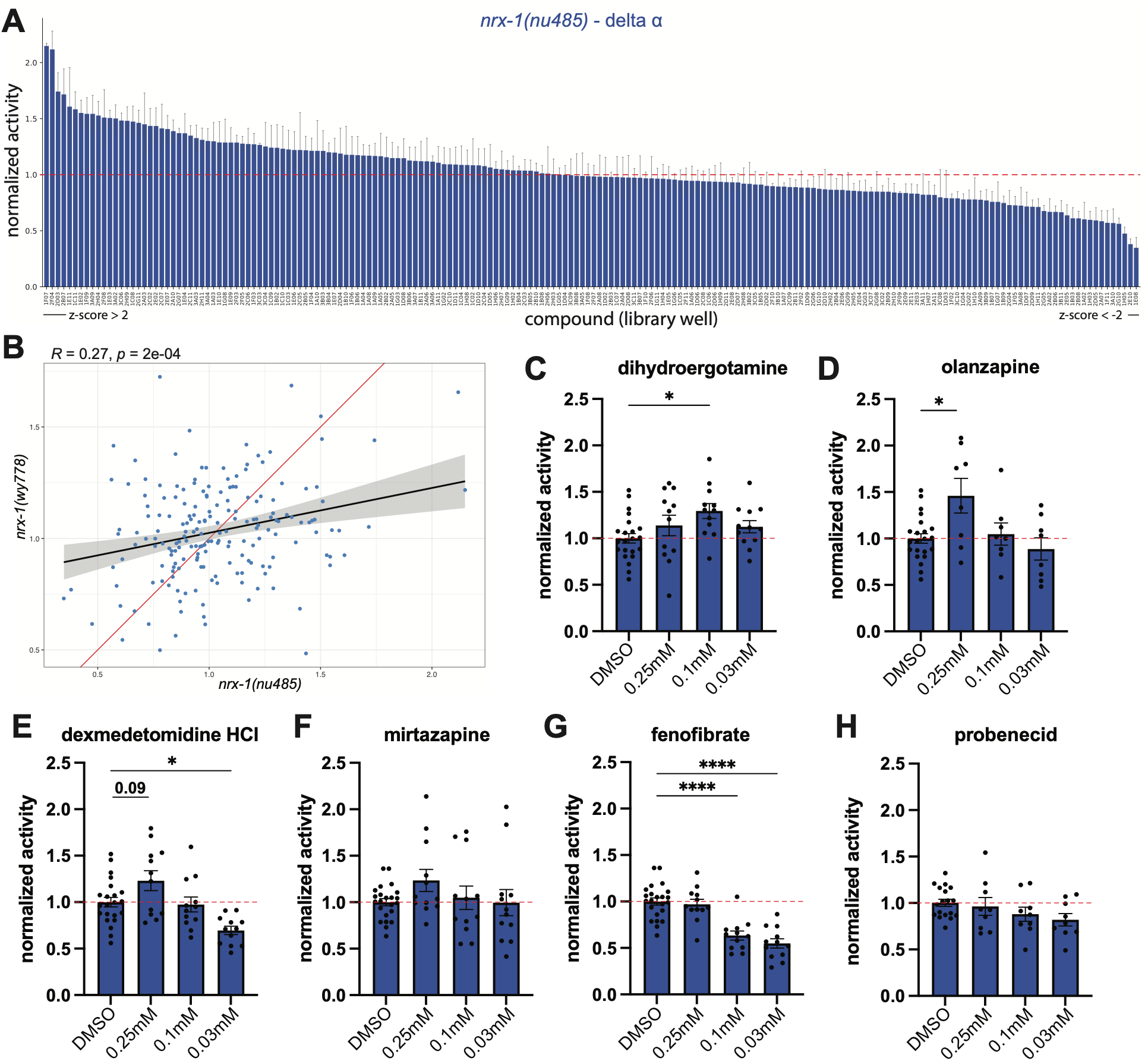
Screen and identification of compounds that increase activity of animals lacking only *nrx-1* α isoform. **A**) Activity levels of ∼30 *nrx-1(nu485)* animals swimming in liquid per well with no food and 0.1 µM of each compound (190 from TocriScreen library) in 1% DMSO normalized to animals in DMSO vehicle alone (1%)(compounds ordered from highest to lowest average normalized activity, black lines under x-axis indicate the compounds with z-scores >2 or <-2 and top 10 compounds further validated in panels below). (**B**) Plot of results from *nrx-1(wy778)* and *nrx-1(nu485)* screens with correlation line (black) and 95% confidence intervals shown (grey)(red line is correlation line of 1 for reference). Activity levels of ∼30 *nrx-1(nu485)* animals swimming in liquid per well with no food and indicated concentrations in 1% DMSO normalized to animals in DMSO vehicle alone for (**C**) Dihydroergotamine mesylate, (**D**) olanzapine, (**E**) dexmedetomidine hydrochloride, (**F**) mirtazapine, (**G**) fenofibrate, and (**H**) probenecid.

To characterize the top compounds from the *nrx-1(nu485)* screen, we tested multiple concentrations of a subset of the compounds. We selected a smaller range of concentrations, including a higher concentration of 0.25 mM, based on the results from previous screen validation experiments (**Figure 2** and **Supplemental Figure 1**). Dihydroergotamine mesylate and olanzapine both significantly increased activity levels of *nrx-1(nu485)* animals at 0.25 or 0.1 mM (**Figure 3**). Dexmedetomidine hydrochloride and mirtazapine slightly increased activity of *nrx-1(nu485)* animals compared to DMSO alone at the highest concentrations, but this was not significant (**Figure 3**). However, low concentrations of dexmedetomidine hydrochloride and fenofibrate significantly decreased activity of *nrx-1(nu485)* animals compared to DMSO alone (**Figure 3**), while probenecid had no impact on activity (**Figure 3**). These results confirmed the impact of a subset of the compounds, with variable or opposite impacts observed for some compounds and concentrations from different sources.

### Monoamine targeting compounds increase the activity of food deprived nrx-1 mutants

The top compounds that impacted both *nrx-1* alleles (and some impacting controls) were of interest for further analysis, in particular, compounds targeting monoamine pathways and receptors. To carefully analyze the impact of each compound across and between genotypes, as well as over time, we designed experiments using the same 96-well WormCamp setup and analysis, but instead with a single animal per well. We ran each compound and *nrx-1* allele with control animals, and normalized activity values to the average activity of control DMSO wells for each plate across 8 hours to closely compare across days, plates, and multiple imaging platforms. First, we analyzed average activities across the 8 hours to compare the magnitude of effect of each compound across the *nrx-1* genotypes. We found that both *nrx-1* alleles displayed reduced the activity compared to control animals (swimming in M9 without food in 1% DMSO)(**Figure 4A**). Overall, we found that the impact of each compound replicated in the single animal setups. Olanzapine and dihydroergotamine (2 concentrations) increased activity of both *nrx-1* alleles, while mirtazapine (2 concentrations) increased activity of only *nrx-1(nu485)* animals (**Figure 4A**). A number of the compounds increase the activity of *nrx-1* alleles at or above activity levels of controls with only vehicle (**Figure 4A**), suggesting rescue or further hyperactivity at these concentrations. In control animals, the compounds had variable impacts compared to the previous experiments (**Supplemental Figure 1D**). While dihydroergotamine (2 concentrations) increased activity levels, there was no impact with asenapine, olanzapine, or mirtazapine (**Supplemental Figure 1D**). The reduced impact for compounds in the single animal assays in controls may represent differences in the magnitude of impact of asenapine and olanzapine in controls (**Figure 2**).

**Figure 4.**
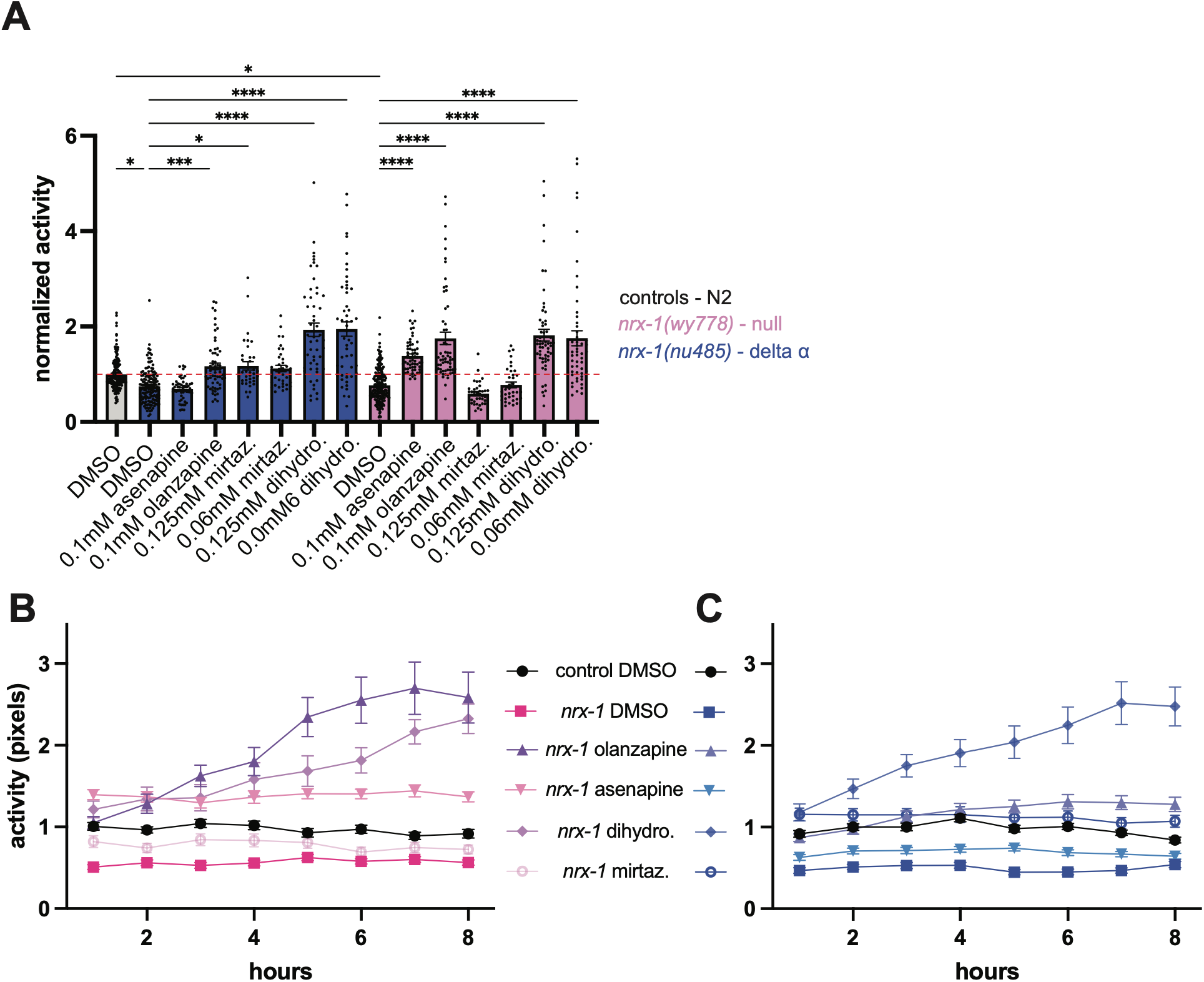
Testing monoamine targeting compounds between genotypes and over time. **A**) Mean activity levels of single animals per well swimming in liquid with no food and 0.1 µM of each compound in 1% DMSO normalized to animals in DMSO vehicle alone (1%) -controls (grey), *nrx-1(nu485)*(blue), *nrx-1(wy778)*(pink). Mean activity levels of single animals per well binned per hour for 8 hours of acute treatment with each indicated compound at 0.1 or 0.125 mM for (**B**) *nrx-1(wy778)*(pink) or (**C**) *nrx-1(nu485)*(blue), controls with DMSO alone for comparison (black).

We also analyzed the temporal impact of the acute treatment of animals with each compound and plotted the single animal assay data over time, focusing on single concentrations (**Figure 4B-C**). Treatment with dihydroergotamine showed increases in activity at even the start of the experiment for both *nrx-1* alleles, with steadily increasing activity across nearly all 8 hours, having a ramping impact on activity (**Figure 4B**). In contrast, olanzapine increased activity of both *nrx-1* alleles with quite different trajectories. In *nrx-1(wy778)* animals, olanzapine treatment resulted in an early increase with steadily increasing activity over the 8 hours (**Figure 4B**), like dihydroergotamine. However, *nrx-1(nu485)* animals responded to olanzapine with an early increase in activity that was maintained but did not further increase over the 8 hours (**Figure 4C**). A similar maintained but not changing increase in activity was observed with asenapine maleate treatment of *nrx-1(wy778)* and mirtazapine treatment of *nrx-1(nu485)*(**Figure 4B-C**). Therefore, the compounds have a distinct impact on activity levels over time that is compound and genotype-specific. Overall, we replicate the impact of most monoamine modifiers across genotypes and find differences in the timing and magnitude of impact between *nrx-1* alleles. Dihydroergotamine drastically increased activity in all genotypes in the single animal assay. While olanzapine is a multi-receptor antagonist that removes inhibition by blocking receptors, dihydroergotamine is a partial agonist (Dopamine, adrenergic, and 5-HT1/2) with broad activation of many receptors, including some auto receptors. Both likely enhance net monoaminergic signaling through disinhibition or direct activation, which can compensate for loss of neurexin by increasing broad monoaminergic tone.

### Differential impact of monoamine targeting compounds on social feeding behavioral phenotypes of nrx-1 alleles

We next wanted to test the impact of the monoamine targeting compounds on other behaviors and circuits. Wild isolate strains of *C. elegans* and other *Caenorhabditis* species display a range of solitary and social feeding behaviors, where social strains aggregate together in clumps in the presence of food and solitary strains actively avoid other animals^39–43^. This behavior is driven by the integration of environmental (oxygen, carbon dioxide, chemosensory, and food) and social (pheromones, touch, and population density) sensory cues. While the control strain N2 Bristol is completely solitary under normal conditions, loss of function variants in the conserved neuropeptide receptor, *npr-1*, induce strong social feeding behavior and serve as a genetically matched social feeding control strain to assay social feeding behavior. We previously reported that loss of function alleles of *nrx-1* (including both *wy778* and *nu485*) significantly reduce the number of aggregating animals, and that *nrx-1* functions in this behavior through modification of specific glutamate sensory neuron synapses (ASH neuron pair)^26^. Social feeding behavioral phenotypes of *nrx-1* alleles can therefore serve as a second functional readout for *nrx-1* isoforms acting at least in part in distinct neurons and circuits.

We first tested the DMSO vehicle and monoamine modifiers on the social control background (*npr-1(ad609))* with and without *nrx-1(wy778)*, and found that DMSO did not impact aggregation and social feeding behavior in either strain. We also confirmed the reduction in the number of aggregating animals with loss of *nrx-1* isoforms (**Supplemental Figure 2**). In social feeding controls, olanzapine, dihydroergotamine, and mirtazapine had no significant impact on the number of aggregating animals, while asenapine and dexmedetomidine hydrochloride significantly decreased the number of aggregating animals to near solitary levels (**Figure 5**, **Table 1, Supplemental Figure 2**). In *npr-1(ad609); nrx-1(wy778)* animals, olanzapine significantly increased the number of aggregating animals, while asenapine and dexmedetomidine hydrochloride significantly decreased the number of aggregating animals (**Figure 5B&C**). Dihydroergotamine and mirtazapine had no significant impact on the number of aggregating *npr-1(ad609); nrx-1(wy778)* animals (**Figure 5B&C**). In social controls and with *wy778*, mirtazapine had a variable impact on the number of aggregating animals, which trended towards a decrease that was not significant (**Figure 5B&C**).

**Figure 5.**
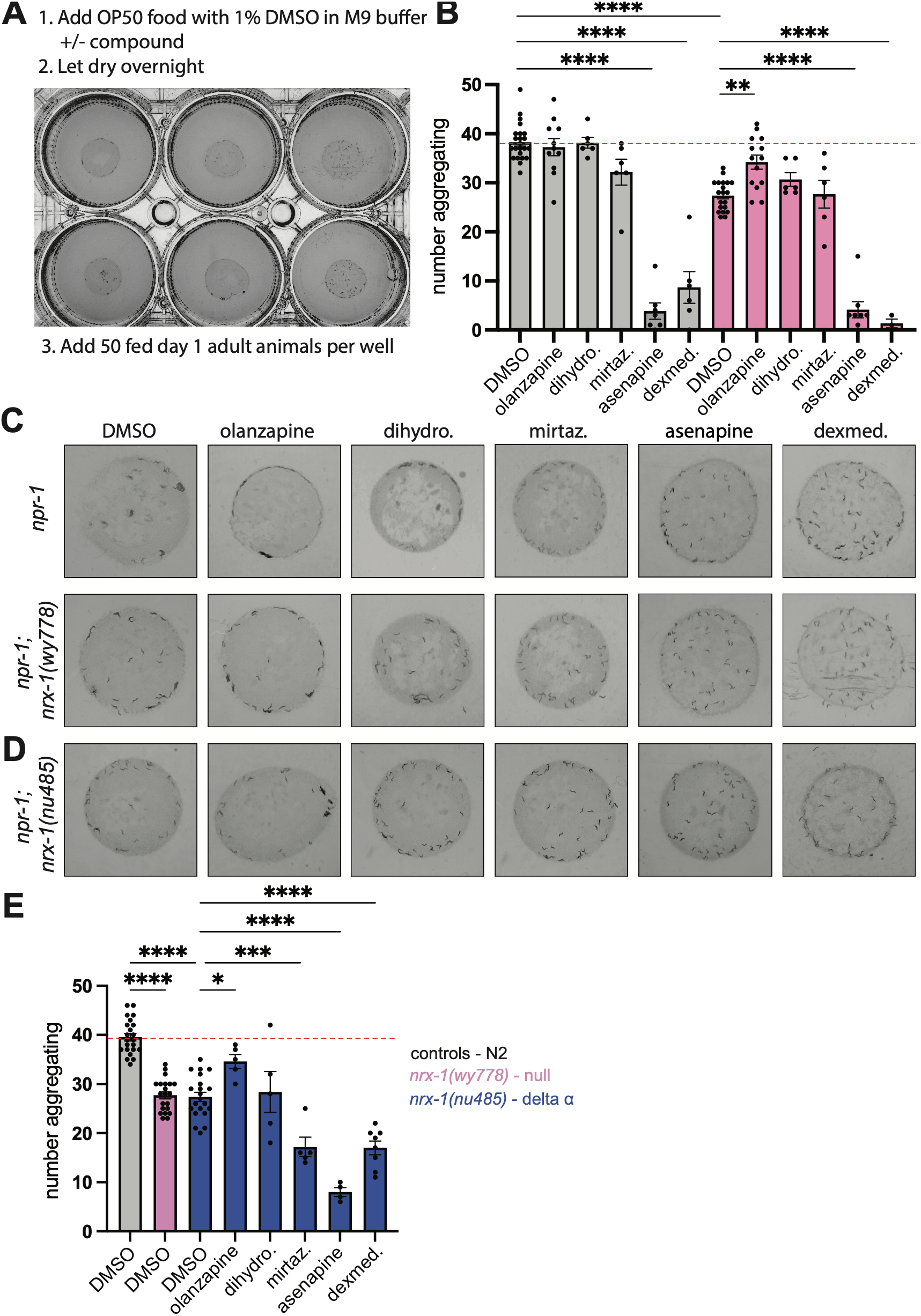
Social feeding behavior and nrx-1 phenotypes are differentially impacted by monoamine targeting compounds. **A)** Image and description of social feeding behavioral assay using 6-well plates. **B)** Quantification and (**C**) representative images of the number of aggregating *npr-1* control (grey) and *npr-1; nrx-1(wy778)*(pink) animals with 1% DMSO in M9 buffer or various compounds as labeled at 0.1mM with 1% DMSO in M9 buffer. **D)** Representative images and (**E**) quantification of the number of aggregating *npr-1* control (grey), *npr-1; nrx-1(wy778)*(pink), or *npr-1; nrx-1(nu485)*(blue) animals with 1% DMSO in M9 buffer or *npr-1; nrx-1(nu485)* animals with various compounds as labeled at 0.1mM with 1% DMSO in M9 buffer.

**Table 1.** Summary of monoamine targeting compounds across behaviors and genotypes. p-values indicated for each compound in controls and nrx-1 allele for each behavioral phenotype (purple increase, green decrease).

| Compound Name | Food deprivation response |  |  |  | Social feeding |  |  |  |
| --- | --- | --- | --- | --- | --- | --- | --- | --- |
|  | controls | *nrx-1(wy778)* | *nrx-1(nu485)* | *nrx-1(sy869)* | controls | *nrx-1(wy778)* | *nrx-1(nu485)* | *nrx-1(sy869)* |
| Olanzapine | 0.9995 | 0.0001 | 0.0006 | 0.0071 | 0.9860 | 0.0100 | 0.0100 | 0.3126 |
| Dihydroergotamine mesylate | 0.0001 | 0.0001 | 0.0001 | 0.0015 | 0.9999 | 0.9999 | 0.9761 | 0.9999 |
| Mirtazapine | 0.9993 | 0.9609 | 0.0125 | 0.0341 | 0.1084 | 0.9558 | 0.0100 | 0.6050 |
| Asenapine maleate | 0.9999 | 0.0001 | 0.9999 | x | 0.0001 | 0.0001 | 0.0001 | 0.8755 |
| *single animal data |  |  |  |  |  |  |  |  |

We next tested the impact of modifiers on the same behavior in *nrx-1(nu485).* We again validated that loss of *nrx-1* α isoform reduced the number of aggregating animals compared to social feeding controls, which was similar to *nrx-1(wy778)*(Figure 5B-D)^26^. Exposure to olanzapine significantly increased the number of *nrx-1(nu485)* aggregating animals, while dihydroergotamine had no significant impact, and mirtazapine, asenapine, and dexmedetomidine hydrochloride reduced social feeding behavior (**Figure 5D&E**). In social feeding behavior, olanzapine increases the number of aggregating animals in both *nrx-1* alleles, suggesting moderate serotonin and adrenergic modulation can compensate for loss of *nrx-1* in this behavioral circuit. We also find strong differences for several compounds between behaviors, with asenapine strongly decreasing social feeding behavior across all genotypes in the social feeding background (*npr-1(ad609)*), and mirtazapine decreasing social feeding specifically in *nu485*, the allele it specifically increased activity in with food deprivation.

### Monoamine targeting compounds improve behavioral phenotypes of a conserved NRXN1/nrx-1 patient variant (L18Q/L16Q)

To assess whether compounds identified using deletion alleles could also impact phenotypes of naturally occurring variants, we tested the top compounds against a conserved missense variant identified in an autism proband^44^. This variant alters the signal sequence of the long α isoform, potentially causing protein misfolding or trafficking defects distinct from simple loss-of-function, and we recently reported that insertion of this variant (L16Q in *C. elegans*, corresponding to the patient-identified L18Q in human *NRXN1* (**Figure 1A**)) into the endogenous

*C. elegans nrx-1* gene results in multiple behavioral phenotypes^44,45^. First, we tested the homozygous *nrx-1(sy869)* allele in the single animal liquid WormCamp with DMSO vehicle alone, and found that *nrx-1(sy869)* animals had reduced activity swimming in liquid with 1% DMSO compared to controls, replicating our previously reported phenotype in liquid culture (**Figure 6A**)^45^. We then tested the impact of two concentrations of top candidate compounds, olanzapine, dihydroergotamine, and mirtazapine, for impact on reduced swimming activity of *nrx-1(sy869)* animals. We find that the lower concentration of olanzapine significantly increased activity of *nrx-1(sy869)* animals, but the higher concentration did not, and that both concentrations of dihydroergotamine also significantly increased activity levels of *nrx-1(sy869)* animals (**Figure 6A**). Mirtazapine also significantly increased activity levels of *nrx-1(sy869)* animals at the higher dose (**Figure 6A**). Therefore, we find that the monoamine targeting compounds from the *nrx-1(nu485)* screen can improve behavioral phenotypes of this conserved variant identified in humans, although the magnitude of impact is reduced compared to deletion alleles, suggesting differences in impact from full loss of the alpha isoform alone (n*u485*). This is further highlighted by our findings that the *sy869* allele can also induce gain of function behavioral phenotypes^45^.

**Figure 6.**
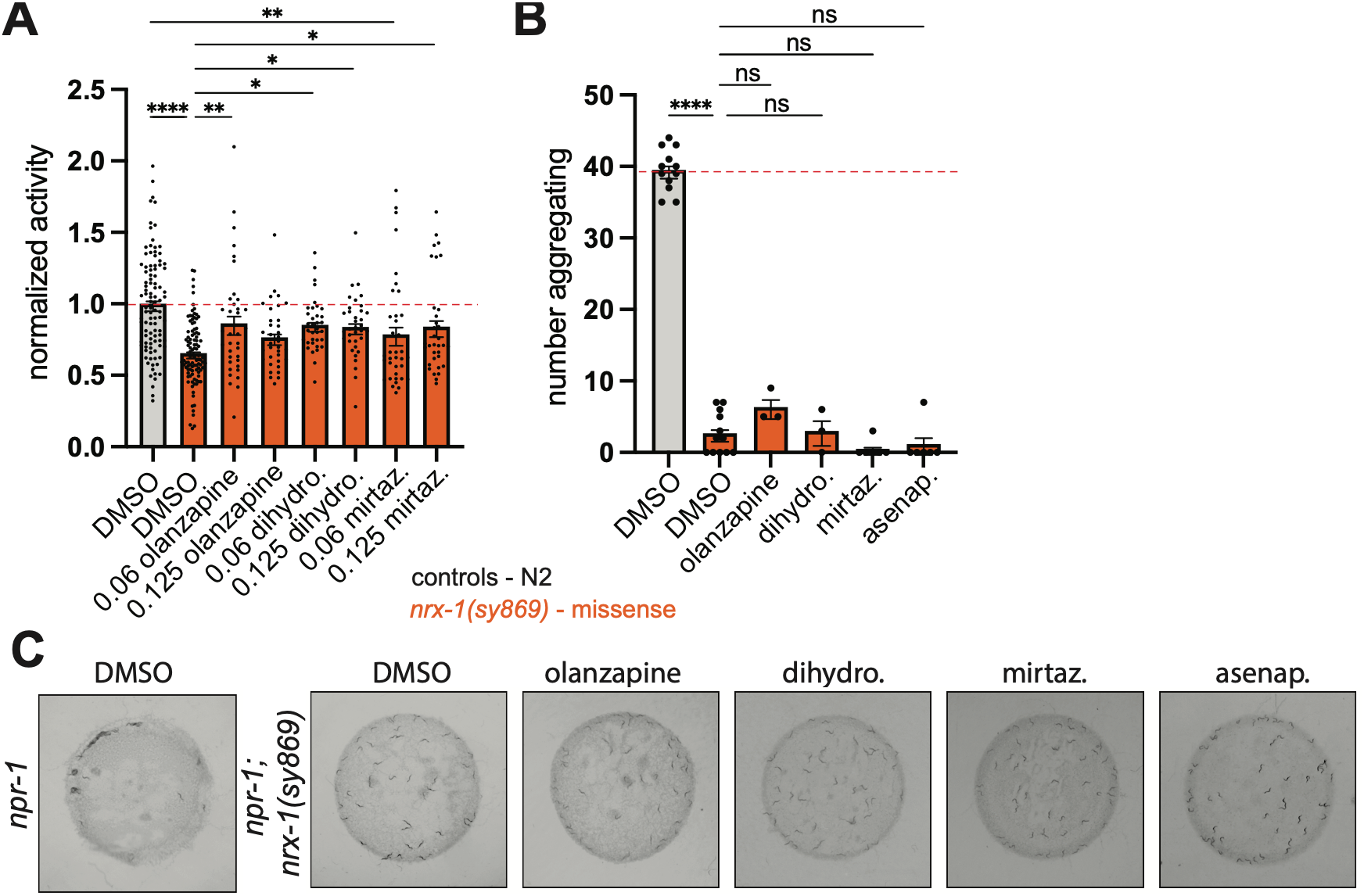
Monoamine targeting compounds restore a behavioral phenotype of a conserved neurexin human variant. **A**) Mean activity levels of single animals per well swimming in liquid with no food and 0.1 µM of each compound in 1% DMSO normalized to animals in DMSO vehicle alone (1%) -controls (grey), *nrx-1(sy869)*(orange). **B)** Representative images and (**C**) quantification of the number of aggregating *npr-1* control and *npr-1; nrx-1(sy869)*(orange) animals with 1% DMSO in M9 buffer or various compounds as labeled at 0.1mM with 1% DMSO in M9 buffer.

Next we tested a subset of these compounds on the reduced social feeding phenotype of this conserved variant^45^. Compared to social control animals, the *npr-1(ad609);nrx-1(sy869)* animals show a strong and significant reduction in the number of animals aggregating, as previously reported (**Figure 6B&C**)^45^. Exposure to olanzapine, dihydroergotamine, mirtazapine, or asenapine had no impact on the number of aggregating *npr-1(ad609);nrx-1(sy869)* animals (**Figure 6B&C**).

### Olanzapine uniquely improves activity and survival phenotypes of NRX-1 in Drosophila

To establish whether monoamine enhancement represents an evolutionarily conserved compensatory mechanism, we tested the leading candidates in *Drosophila melanogaster*, a system where monoamine signaling similarly regulates activity-dependent behaviors. A number of *Drosophila* studies have identified behavioral phenotypes associated with the loss of *Nrx-1*^8,9^, which encodes only a single long α isoform. First, we generated animals with loss of *Nrx-1* by combining two deletion alleles as trans-heterozygotes (**Figure 7A**). Control and *Nrx-1* animals were pre-treated with compounds for 6 hours, then activity with or without compound was monitored for at least 4 days (**Figure 7B**). *Nrx-1* animals are frail, and show dramatically reduced activity levels during both days and nights compared to control animals in the presence of DMSO vehicle alone (**Figure 7C-D**). Exposure to 0.25 mM of olanzapine significantly increased activity levels of *Nrx-1* animals across 0-36 hours compared to *Nrx-1* animals on vehicle control (**Figure 7C**). The impact of olanzapine on activity levels was also observed to a lesser extent in control animals, with the higher concentration of olanzapine significantly increasing activity of control animals at 12-24 hour and 24–36-hour timepoints (**Figure 7C**). We performed the same behavioral activity analysis using asenapine maleate and dihydroergotamine, but these compounds did not increase activity of *Nrx-1* or control animals (**Supplemental Figure 3**).

**Figure 7.**
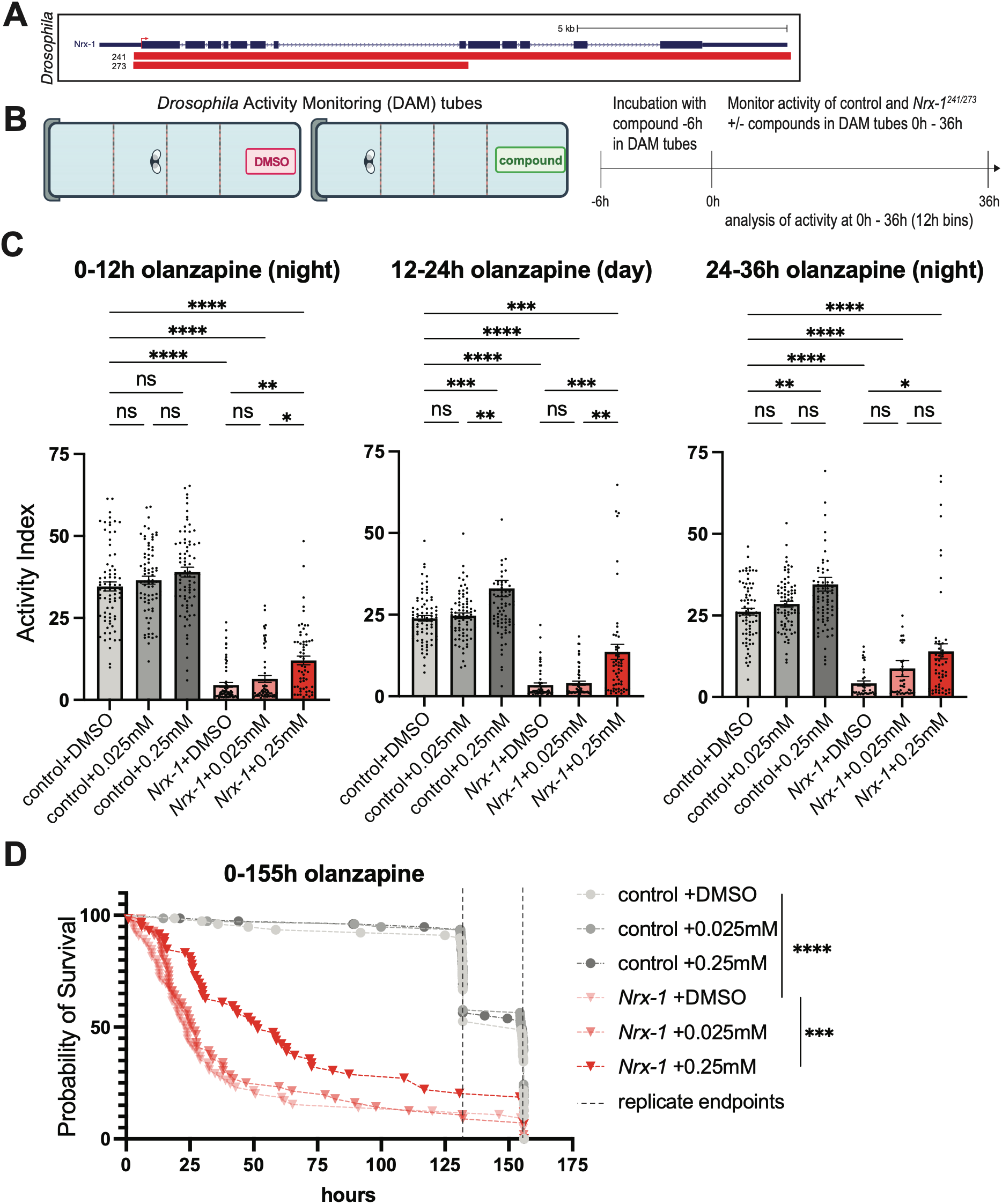
Olanzapine improves reduced activity of *Nrx-1 Drosophila*. **A)** Diagram showing *Nrx-1* genomic locus, with deletion alleles used in this study shown in red. **B)** Cartoon of DAM system for monitoring *Drosophila* behavior and activity and timeline of experiments (3 infrared beams depicted, actual instruments have 15 beams). **C)** Activity levels of Drosophila across day and night, for controls and trans-heterozygous *Nrx-1^273/241^* flies with DMSO or olanzapine at 0.025mM and 0.25mM. **D)** Graph of the probability of survival for controls and trans-heterozygous *Nrx-1^273/241^* flies with DMSO or olanzapine at 0.025mM and 0.25mM (dashed lines represent end of experiment for control flies at either 135 or 155 hours).

In activity assays, we observed that *Nrx-1* animals show a significantly reduced lifespan compared to control animals (**Figure 7D**). To assess whether olanzapine improves this phenotype in addition to activity levels, we examined the lifespan in animals treated with olanzapine compared to vehicle control and found that exposure to the higher concentration of olanzapine extended lifespan in *Nrx-1* flies. We did not observe any impact of olanzapine on survival in control animals for the length of the experiment, which only analyzed lifespan out to 135 or 155 hours (indicated with dashed lines)(**Figure 7D**). Taken together, our cross-species analyses identify monoamine signaling as a conserved pathway capable of modifying neurexin-associated behavioral dysfunction.

## DISCUSSION

### Monoamine enhancement can compensate for neurexin dysfunction

The pharmacological screens here identify monoamine-targeting compounds as modifiers of *nrx-1* behavioral phenotypes, confirming a functional relationship between neurexin and monoamine signaling. This finding is significant given our previous demonstration that *nrx-1* isoforms differentially impact food response through the RIC interneuron circuit, which uses octopamine (the invertebrate analog of norepinephrine)^25^, serotonin, and dopamine. These results extend the relationship, revealing that pharmacological modulation of serotonin, dopamine, and octopamine can bypass or compensate for deficits in synaptic organization or function caused by neurexin loss. Several mechanisms could explain the monoamine-neurexin interaction in the context of the food response circuits (**Figure 8**). First, neurexins may impact the release of monoamines from pre-synaptic sites or monoamine receptors function at post-synaptic sites, similar to established roles at glutamate and GABA synapses^46,47^. Loss of neurexin could reduce monoamine signaling efficiency or receptor availability at specific synapses, which pharmacological enhancement can overcome through compensatory receptors or parallel pathways. Second, monoamine signaling might act in parallel circuits to restore behavioral output despite synaptic deficits in neurexin lacking neurons. Our finding that different compounds work in different *nrx-1* alleles supports the idea that distinct isoforms interface with monoamine systems through separate mechanisms. The temporal dynamics we observed, where some compounds produced immediate effects (asenapine, mirtazapine) while others show increasing impact over time (dihydroergotamine, olanzapine in *wy778*), suggest both acute release or receptor-level modulation and longer-term adaptive changes. This aligns with the known dual actions of monoamines in neuronal activity and slower transcriptional changes^47^. The ramping effect of dihydroergotamine suggests it triggers adaptive responses that progressively enhance circuit function, whereas the rapid plateau effects of asenapine and mirtazapine suggest more direct modulation.

**Figure 8.**
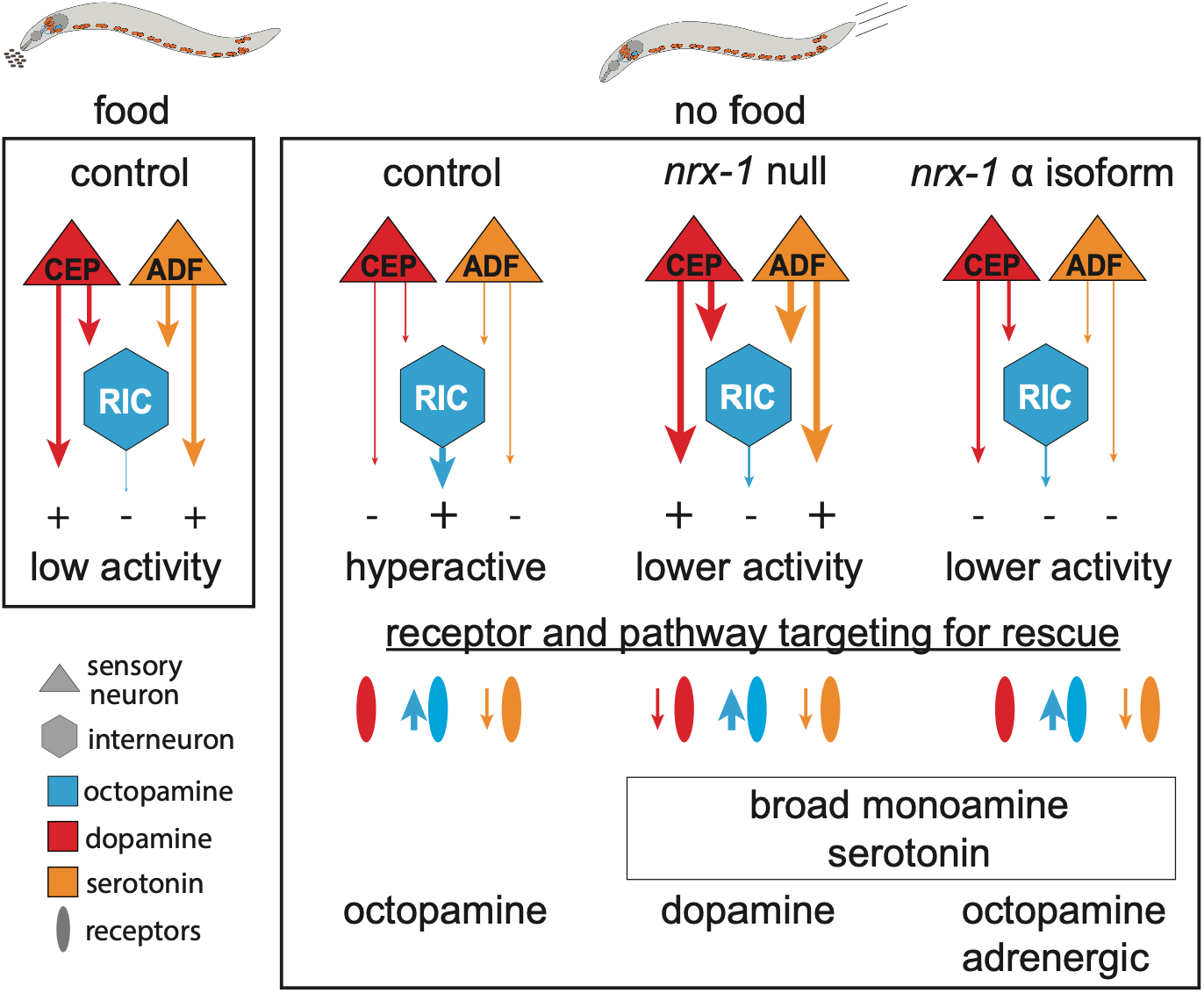
Cartoon of *C. elegans* monoamine signaling in food response behaviors in controls and *nrx-1* alleles. Control animals on food have low activity levels driven by serotonin and dopamine signaling, with reduced octopamine signaling. Control animals without food become hyperactive driven by increase octopamine signaling and reduced serotonin and dopamine signaling. In *nrx-1* null animals, there is decreased octopamine signaling, and unchanged or increased dopamine and serotonin signaling – activity levels can be increased with broad monoamine modulators, or serotonin and dopamine antagonists. In animals lacking *nrx-1* α isoform, there is decreased octopamine response, and mostly unchanged dopamine and serotonin signaling – activity levels can be increased with broad monoamine modulators, exogenous octopamine or adrenergic receptor agonists.

### Isoform-specific pharmacology reveals differential monoamine dependencies

A striking finding from our study is the differential pharmacological response of the two loss of function *nrx-1* alleles and the L16Q missense *nrx-1* allele. While olanzapine increased activity in all three alleles, other compounds showed remarkable specificity, indicating specific circuit disruptions that likely represent isoform-specific mechanisms. Neurexin loss causes increased serotonin signaling or sensitization as behavioral rescue required blocking postsynaptic serotonin receptors (olanzapine (5-HT2A antagonist), asenapine (5-HT2A/2C antagonist), mirtazapine (5-HT2A/2C/3 antagonist), dihydroergotamine (5-HT antagonist)), while activating them had the opposite impact (vilazodone (5-HT1A agonist)(**Figure 8**).

The selective efficacy of adrenergic modifiers in *nu485* mutants suggests that loss of α isoform may decrease octopaminergic/adrenergic tone or create a disruption which can be compensated by enhanced adrenergic signaling (**Figure 8**). Mirtazapine (α2 antagonist), dexmedetomidine (α2 agonist), and dihydroergotamine (α partial agonist), possess dopaminergic activity and rescue *nu485* while showing lesser impact in null mutants. This distinguishes them mechanistically from the broader multi-target modifiers effective in both alleles and suggests a distinct mechanistic role for the γ isoform in responding to adrenergic/octopaminergic signals. The efficacy of both α2 agonists and antagonists likely reflects differential localization and function of α2 receptors. Presynaptic α2 auto receptors provide negative feedback, reducing release when activated; their blockade by mirtazapine could therefore enhance adrenergic tone. Postsynaptic α2 receptors mediate direct signaling effects; their activation by dexmedetomidine directly stimulates downstream pathways. Enhanced adrenergic signaling can therefore increase activity through increased transmitter release (α2 antagonism) or direct receptor activation (α2 agonism)(**Figure 8**). This would predict that α isoform *nrx-1* alleles would respond to increase adrenergic tone via exogenous octopamine, while nulls would not, which is consistent with our previous findings^25^, although this may differ in missense *nrx-1* alleles that have complex impact on function of isoforms.

In contrast, the response of *wy778* mutants (lacking both α and γ isoforms) to asenapine, pimethixene maleate, and perospirone, which have strong dopamine (D2) antagonist activity in addition to serotonin effects, suggests that complete neurexin loss requires broader monoamine modulation and dopaminergic signaling (**Figure 8**). Minimal rescue was observed with compounds targeting only D2 receptors or dopamine signaling in either allele. The observation that mirtazapine, which lacks any dopaminergic activity, shows no efficacy in *wy778*, but significant efficacy in *nu485* provides quantitative support for isoform-specific mechanistic divergence. Loss of the γ isoform appears to increase dopaminergic tone or creates a disruption that can be rescued by blocking dopamine signaling (**Figure 8**).

Lastly, olanzapine’s efficacy across all three alleles can be understood through its multi-receptor profile. As a moderate antagonist at dopamine D2, serotonin 5-HT2A, α1-adrenergic, and histamine H1 receptors, olanzapine modulates multiple monoamine systems to compensate for isoform-specific (*nu485*) and complete neurexin (*wy778*) loss or disruption (*sy869*)(**Figure 8**), further demonstrating neurexin isoform-specific roles across monoamine signaling pathways. *NRXN1* variants often affect specific exons and isoforms^20,22,2,19,4,49^, and our results suggest that interventions need to be tailored based on the isoforms disrupted. α isoform disruptions might respond to adrenergic agents, while those affecting multiple isoforms might require broader monoamine modulation.

### Behavior-specific compound effects reflect circuit-level organization

Our multi-behavior approach revealed pharmacological complexity not evident in single-behavior screens and identified novel roles for monoamine signaling in *C. elegans* social feeding behavior. While olanzapine improved both food deprivation responses and social feeding behaviors in *nrx-1* loss of function alleles (but not social feeding of the *sy869* missense allele), asenapine and mirtazapine showed striking behavioral divergence: enhancing locomotor responses to food deprivation but eliminating social feeding behavior (even in control animals). This behavior-specific pharmacology reflects the distinct circuits underlying these behaviors. Food deprivation response requires a balance of octopamine signaling from RIC interneurons and integration with dopamine, serotonin and other pathways (**Figure 8**), whereas social feeding depends on glutamate and acetylcholine signaling from sensory neurons with integration through RMG interneurons and neuropeptide signaling^32,31,25,41,26^. Asenapine’s antagonism of multiple serotonin receptors (5-HT2A, 5-HT2C, 5-HT7) may enhance activity-promoting pathways while simultaneously disrupting the sensory integration required for social feeding. The compound’s particularly potent 5-HT7 antagonism, a property not shared by olanzapine, may disrupt the social feeding circuit.

These findings have significant implications for understanding and treating behavioral heterogeneity in autism and neurodevelopmental disorders involving neurexin and related genes. Complex behavioral profiles across domains including social communication, repetitive behaviors, anxiety, depression, and seizures, where interventions improving one domain may worsen others^43^. Asenapine’s rescue of one circuit-dependent behavior while eliminating another, provides a gene-specific mechanism for behavioral distinctions, underscoring the need to test modifiers across multiple relevant behavioral domains. The identification of olanzapine as uniquely effective across both behaviors suggests that moderate, multi-target receptor modulation is preferable to selective targeting when treating complex behavioral phenotypes spanning multiple neural circuits.

### Comparison to previous behavioral *C. elegans* small molecule modifier screens

Previous *C. elegans* pharmacological screening across four conserved autism-associated gene mutants (*unc-44(ANK2)*, *egl-19(CACNA1C)*, *nmr-2(GRIN2B)*, and *unc-2(CACNA1A*)) found minimal pharmacological convergence, with each gene showing distinct modifier compound profiles^50^. Our observation of weak correlation between modifier compounds for alleles targeting the two neurexin isoforms extends this finding, that even within a single gene, isoform-specific deletions require distinct modifiers. The predominance of monoamine-targeting compounds in our neurexin screens contrasts markedly with the broad-spectrum anticholinergic compounds identified across multiple autism gene mutants (representing 14-20% of hits across *unc-44, egl-19, nmr-2*, and *unc-2* mutants)^50^. This may reflect general motor circuit disruption and compensation or correction of excitatory/inhibitory imbalances. These differential findings suggest specific organizational roles for neurexin isoforms at monoamine synapses, as demonstrated by our previous work. Notably, the *nmr-2(GRIN2B)* mutant also responded to monoamine modulators (metergoline, tryptoline), suggesting that synaptic organization genes (adhesion (neurexin) or signaling (NMDA receptors)) may have distinct vulnerabilities from general motor deficits observed in scaffold genes like *unc-44/ANK2*. Disruption of multiple synaptic organizing molecules creates monoamine dependencies which may represent a broadly applicable compensatory mechanism. The efficacy of monoamine enhancement may reflect neuromodulatory compensation of different circuit dysfunctions, a mechanism that could be broadly applicable across a multitude of genetic and behavioral etiologies.

### Cross-species validation and evolutionary conservation

Our demonstration that olanzapine, but not asenapine or dihydroergotamine, can improve a *Drosophila Nrx-1* behavioral phenotype provides essential evidence for evolutionary conservation of the neurexin-monoamine relationship and demonstrates potential specific mechanistic targeting. *Drosophila* also rely on octopamine signaling, and a balance of monoamines in modifying activity and responses to food and stressors, including food deprivation^51,52^. This evolutionary conservation, spanning over 800 million years from nematodes to insects, substantially strengthens monoamine enhancement as a potential intervention in mammalian and human neurexin models. Our findings have practical implications for prioritizing compounds for mammalian validation; moderate multi-target modulators may have higher translational probability than much more highly selective modifiers. The cross-species conservation of olanzapine’s impact is interesting given that *Drosophila Nrx-1* contains only α-like isoforms, lacking the β and γ isoforms present in mammals and the γ isoform present in *C. elegans*^53^. Since olanzapine improves behavioral phenotypes in both *C. elegans* with α-deletion and *Drosophila Nrx-1* mutants (which only have α), this provides evidence that the α isoform-monoamine interface is an evolutionarily conserved target. Mammalian *NRXN1* deletions and variants affecting α isoforms may therefore be most amenable to monoamine-based interventions. A study using a drug-gene interaction database to associate promising compounds with insomnia risk genes, including *NRXN1*, found that duloxetine hydrochloride and nicotine polacrilex were potential compounds for targeting *NRXN1* associated insomnia^54^. Notably, we found that duloxetine hydrochloride, a serotonin-norepinephrine reuptake inhibitor, increased activity in the *nrx-1(nu485)* screen (ranked 15, z-score 1.5543), providing some experimental support for an interaction of this compound with neurexin.

### Patient variant validation provides clinical relevance and identifies potential complications

While deletion variants represent severe disruptions of neurexin function, most human *NRXN1* variants are missense changes or partial deletions likely impacting specific isoforms. Our demonstration that compounds effective in deletion mutants can also rescue behavioral phenotypes of the conserved neurexin L16Q patient variant (L18Q in human), provides important translational relevance. However, the reduced magnitude of rescue in the missense variant compared to deletion alleles may reflect the distinct mechanistic consequences of protein truncation versus altered protein sequence. This has important implications: certain variants may respond differently to monoamine modulators depending on whether they cause loss-of-function, partial loss-of-function, gain-of-function, or altered protein trafficking. Future studies should include screening for modifiers of this specific allele and consider variant-specific responses and the need for precision medicine approaches beyond gene-level stratification. Our use of the missense variant serves as further proof-of-principle that human patient variants can be functionally modeled and studied using invertebrate organisms.

### Broader implications for synaptic adhesion molecule disorders and towards genotype-guided therapy

Our previous work demonstrated that *C. elegans nlg-1* and *nlr-1* (*CNTNAP* ortholog) mutants have behavioral changes amenable to pharmacological screening^26,27^, and preliminary evidence shows that monoamine modulators also impact behaviors in these genetic backgrounds, but not in all genetic backgrounds. The mechanistic diversity revealed by comparing our results to other autism gene screens has important implications for future intervention approaches^48^. *egl-19/CACNA1C* responded to ROS-modulating polyphenols, reflecting calcium channel roles in oxidative signaling; *nmr-2/GRIN2B* benefited from monoamine modulators and memory enhancers, consistent with functions in synaptic plasticity; and *unc-44/ANK2* showed structural defects amenable to compensation with anticholinergic compounds. This mechanistic heterogeneity matches the functionally diverse categories of conserved autism-associated genes that somehow converge on behavioral phenotypes. This argues for mechanism-based intervention and gene/variant stratification, considering the heterogeneity of behavioral conditions while providing a framework for the development of interventions based on underlying molecular pathways. Synaptic organizing molecules like neurexin function during development and throughout life to coordinate the assembly and maintenance of synaptic molecular complexes. When these organizers are lost, the synaptic machinery may remain largely intact but disorganized or inefficiently deployed. Pharmacological enhancement of signaling through this machinery, particularly through neuromodulatory systems that broadly influence circuit function, may compensate for organizational deficits through many mechanisms (increasing signal-to-noise ratios or recruiting additional synaptic resources).

The identification of olanzapine as effective across neurexin isoform deletions and a conserved missense allele, multiple behaviors, and conserved across species, provides compelling evidence for further translational research. Olanzapine is an FDA-approved atypical antipsychotic already prescribed for schizophrenia and bipolar disorder, conditions also associated with *NRXN1*^20,4,6^. Our findings provide a mechanistic rationale for olanzapine effectiveness and suggest it should be investigated for *NRXN1* deletion carriers. Olanzapine’s multi-receptor and signaling pathway profile appears suited to compensate for both isoform-specific and broader neurexin dysfunction. A retrospective clinical database study to identify individuals with *NRXN1* variants and compare treatments and responses between *NRXN1* carriers and controls prescribed olanzapine versus other antipsychotics. This could justify prospective study of olanzapine treatment in *NRXN1* carriers. Our finding of isoform-specific modifiers also has clinical implications because human *NRXN1* deletions differentially impact exons and disrupt a subset of isoforms. Our finding that mirtazapine specifically improves α-isoform deletion (*nu485*), with no efficacy in the null deletion (*wy778*), while other compounds show opposite patterns, suggests that intervention should be guided by which exons are deleted and isoforms impacted. This represents isoform-level pharmacogenomics, where selection of therapeutic interventions rarely considers specific variants.

### Limitations and future directions

Several important questions remain for future investigation. First, we do not yet understand the molecular mechanisms by which monoamine enhancement compensates for neurexin loss. Do these modifiers act in the same neurons where neurexin normally functions, or through parallel compensatory circuits? Does monoamine modulation restore disrupted receptor trafficking and synaptic organization, or does it bypass structural deficits to improve behavioral output through alternative signaling? Second, our temporal analysis revealed different response dynamics for different compounds, but we tested only acute exposure (4-8 hours). Chronic treatment might reveal different efficacy profiles, tolerance, or adaptive responses. Additionally, our studies in *C. elegans* focused on adult animals; whether developmental exposure would show different effects or alter phenotype emergence remains unexplored. Third, while our cross-species validation in *Drosophila* is valuable, vertebrate and mammalian validation is essential. The substantially greater complexity of mammalian nervous systems, the presence of three neurexin genes (*NRXN1-3*) with extensive alternative splicing, and the different behavioral repertoires all mean that findings may not translate. Fourth, we have not yet identified which specific monoamine receptors mediate the modifications observed, or whether the modifier receptor targeting profiles and specificities are conserved to *C. elegans* receptors. While our isoform-specific compound patterns strongly implicate adrenergic/octopaminergic receptors for α-isoform rescue and broader monoamine modulation for complete nulls, direct evidence requires testing compounds in receptor mutant backgrounds (*ser-1, ser-4, dop-1, dop-2*, etc.). Fifth, our behavior-specific effects underscore the importance of testing across multiple relevant behavioral domains across species, olanzapine has well-described negative impacts in humans, so this question will require systematic investigation. Finally, while we focused on neurexin, the principles we have established, that synaptic molecules loss creates behavioral changes amenable to pharmacological compensation, likely applies broadly to other genes.

Future work should focus on: (1) identifying the specific monoamine receptor subtypes responsible for behavioral rescue, (2) determining whether compounds act in the same neurons where neurexin functions or through parallel circuits, (3) testing whether behavioral improvements correlate with restoration of synaptic structure or function, (4) expanding to screen larger compound libraries and behavioral phenotypes of additional genes, and (5) continuing translation to vertebrate and mammalian models.

---

**Supplemental Figure 1.**
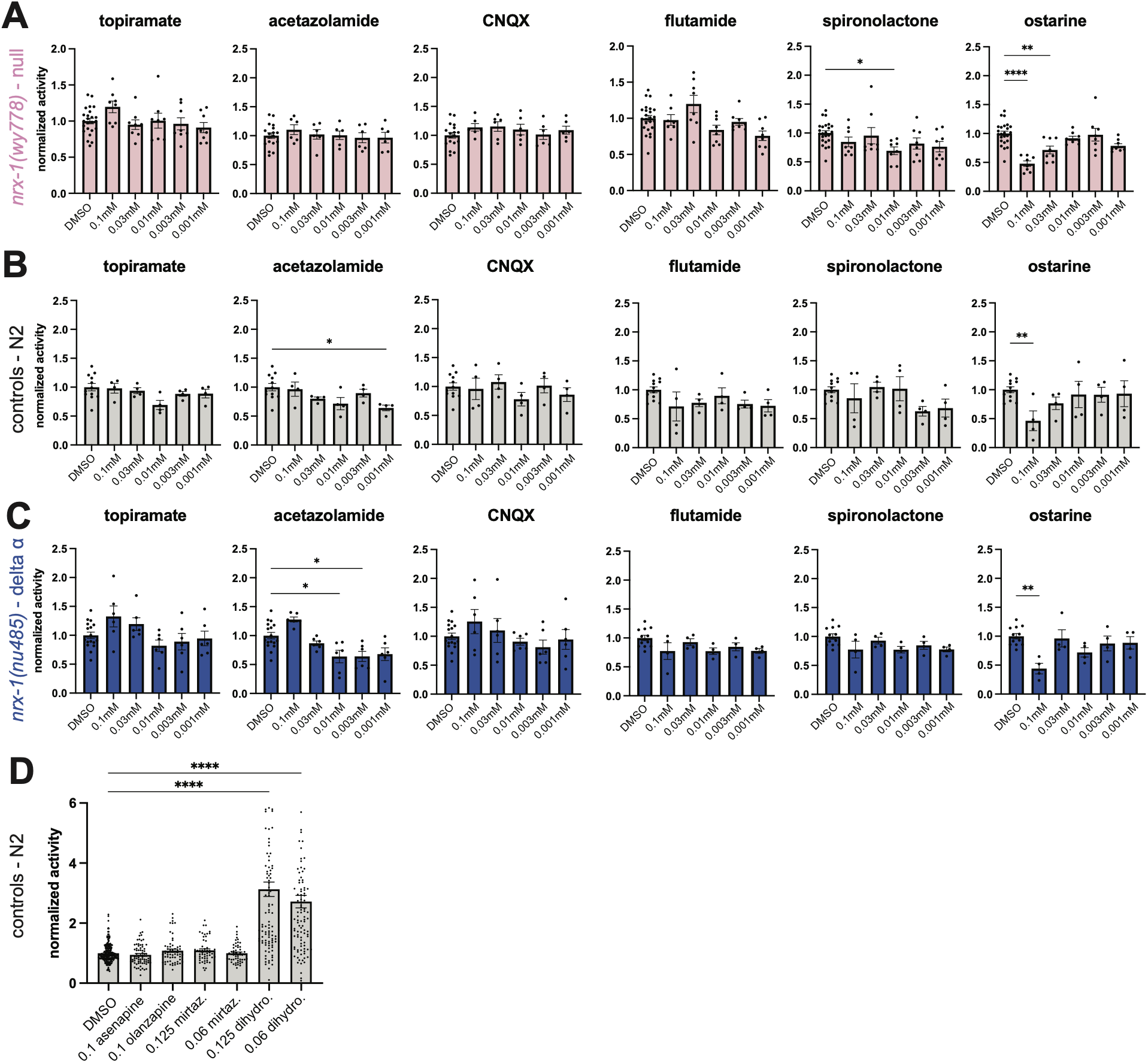
Analyzing compounds identified in null *nrx-1* screen with limited validation. Activity levels of ∼30 animals per well swimming in liquid with no food and a range of concentrations of each compound with 1% DMSO normalized to animals in DMSO vehicle alone (1%) for (**A**) *nrx-1(wy778)*(pink), (**B**) N2 controls (grey), and (**C**) *nrx-1(nu485)*(blue). **D**) Mean activity levels of single N2 control animals swimming in liquid per well with no food and 0.1 µM of each compound in 1% DMSO normalized to animals in DMSO vehicle alone (1%).

**Supplemental Figure 2.**
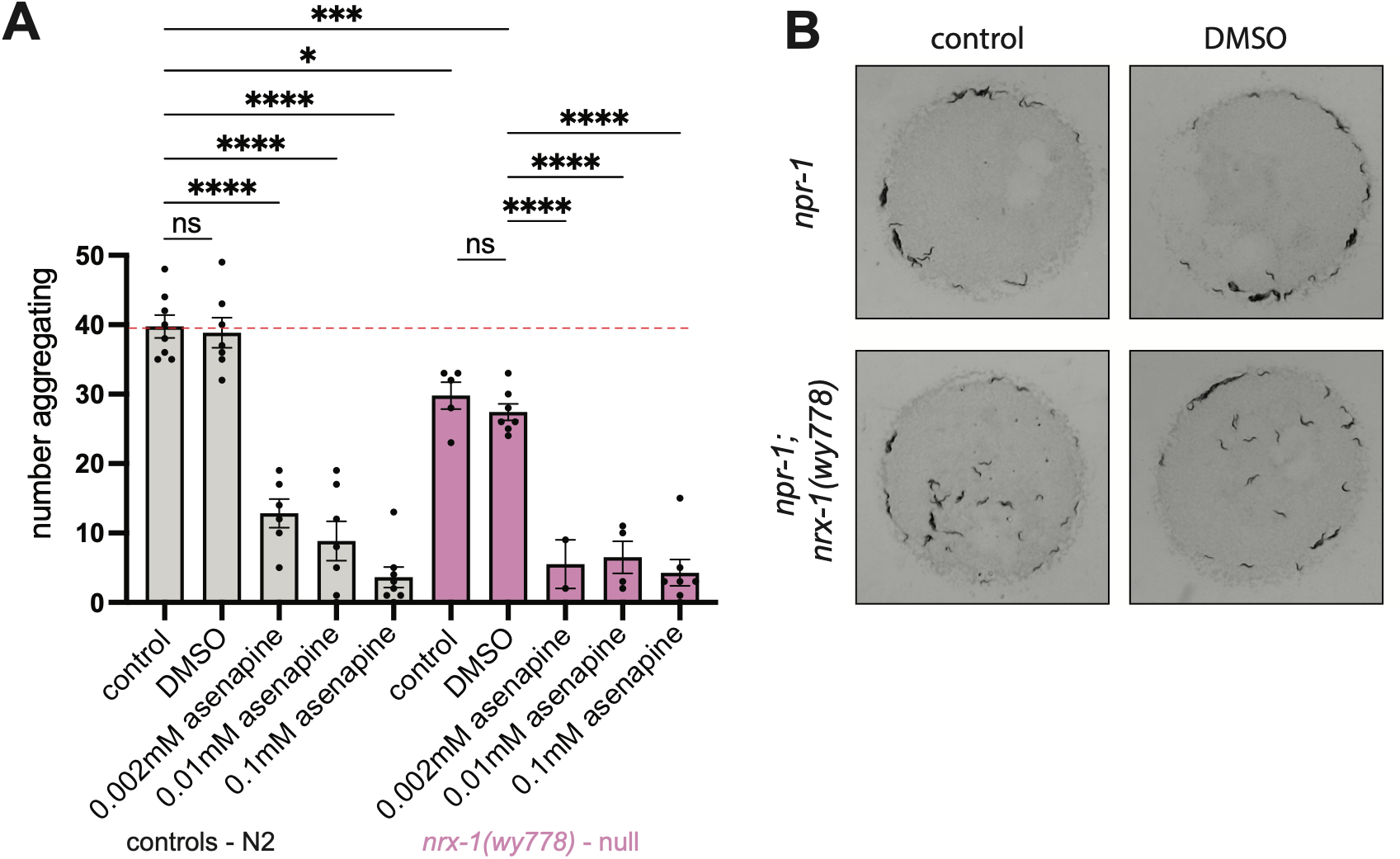
Social feeding behavioral phenotypes in *nrx-1* alleles. **A)** Quantification of the number of aggregating *npr-1* control and *npr-1; nrx-1(wy778)* animals with just OP50 food, 1% DMSO in M9 buffer, or asenapine at multiple concentrations with 1% DMSO in M9. **B)** Representative images of *npr-1* control and *npr-1; nrx-1(wy778)* animals with just OP50 food or 1% DMSO in M9 buffer.

**Supplemental Figure 3.**
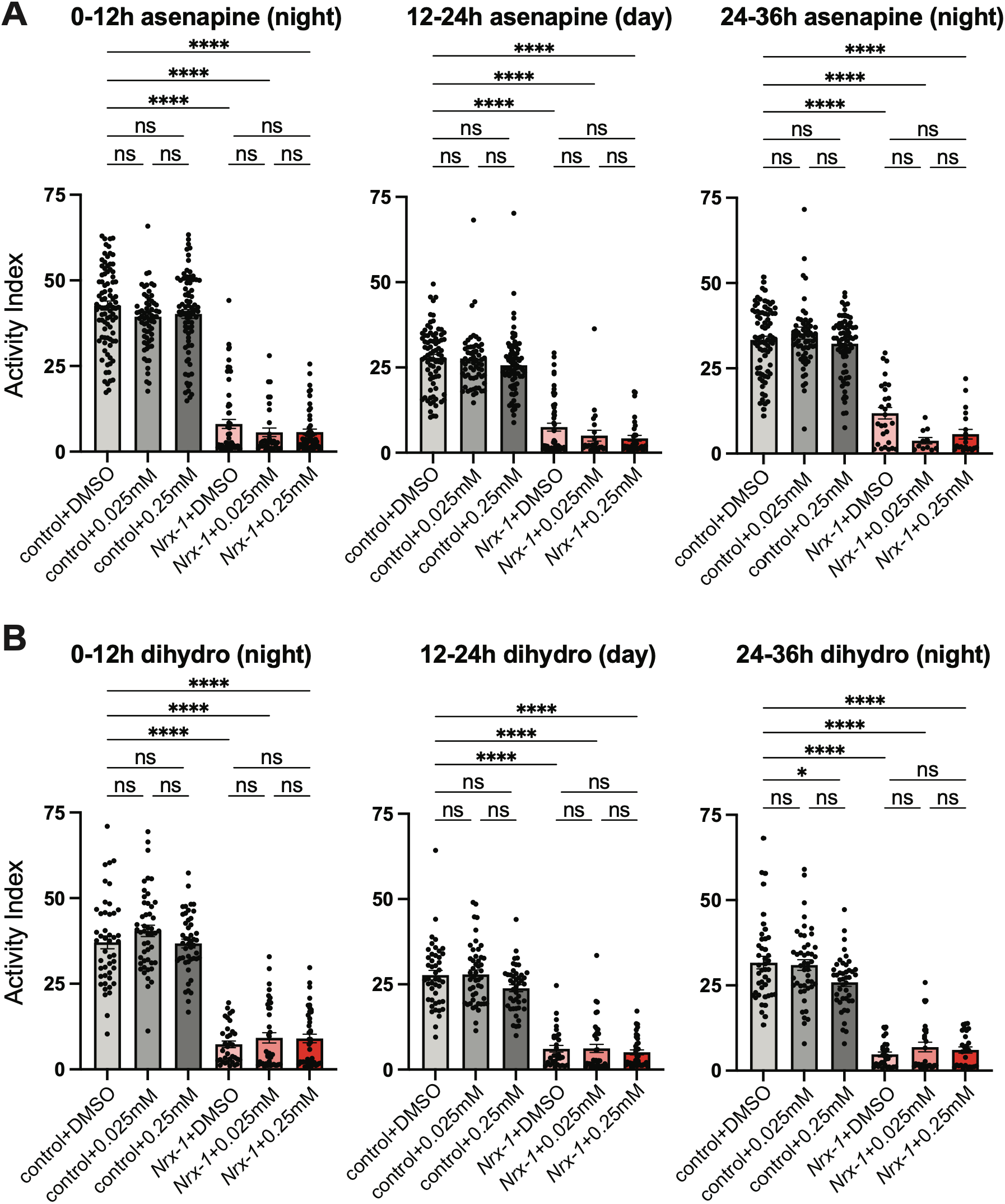
Asenapine maleate and dihydroergotamine do not improve activity phenotypes of *Nrx-1 Drosophila*. **A)** Activity levels of Drosophila across day and night, for controls and trans-heterozygous 273/241 flies with DMSO or asenapine maleate at 0.025 mM and 0.25 mM. **B)** Activity levels of Drosophila across day and night, for controls and trans-heterozygous 273/241 flies with DMSO or dihydroergotamine at 0.025 mM and 0.25 mM.

**Supplemental Table 1.** Small molecule screening results in *nrx-1(wy778)*.

**Supplemental Table 2.** Second source small molecule compound validation.

**Supplemental Table 3.** Small molecule screening results in *nrx-1(nu485).*

**Supplemental Table 4.** Comparison of Z scores from *nrx-1* behavioral screens.

**Supplemental Table 5.** *C. elegans* strains used in this study.

## Acknowledgements

The authors thank the members of the Hart, Kayser, and Jongens labs for technical assistance. The authors thank all members of the Autism Spectrum Program of Excellence, especially Maria Fasolino, and Anthony D. Fouad (Tau Scientific) for technical support. We thank the UPenn High-throughput screening core, RRID: SCD_022379, for their assistance in generating compound concentration gradient plates. Some strains were provided by the CGC, which is funded by NIH Office of Research Infrastructure Programs (P40 OD010440). This work was supported by the Autism Spectrum Program of Excellence and by NIGMS of the NIH under 1R35GM146782 (MPH).

## Author Contributions

MPH conceived and designed the study and experiments with input from TAJ, MSK, and BLB. WS, WH, and RK conducted the behavioral screening and validation experiments with assistance from BLB. WS, JP, and RK performed social feeding behavioral experiments. JNB and MP performed Drosophila assays. MPH processed and analyzed all *C. elegans* data with assistance from WS, WH, and MP, and MP processed and analyzed Drosophila data with assistance from JNB and MSK. MPH wrote the manuscript and compiled the graphs and figures with assistance from MP, WS, JP, JNB, and MSK. All authors reviewed, revised, and approved the manuscript.

